# SPARTA: Interpretable functional classification of microbiomes and detection of hidden cumulative effects

**DOI:** 10.1101/2024.03.07.583888

**Authors:** Baptiste Ruiz, Arnaud Belcour, Samuel Blanquart, Sylvie Buffet-Bataillon, Isabelle Le Hüerou-Luron, Anne Siegel, Yann Le Cunff

**Affiliations:** Univ. Rennes, Inria, CNRS, IRISA, Rennes, France; Univ. Grenoble Alpes, Inria, Grenoble, France; Institut NuMeCan, INRAE, INSERM, Univ Rennes, Saint-Gilles, France; Department of Clinical Microbiology, CHU Rennes, Rennes, France

## Abstract

The composition of the gut microbiota is a known factor in various diseases, and has proven to be a strong basis for automatic classification of disease state. A need for a better understanding of this community on the functional scale has since been voiced, as it would enhance these approaches’ biological interpretability. In this paper, we have developed a computational pipeline for integrating the functional annotation of the gut microbiota to an automatic classification process, and facilitating downstream interpretation of its results. The process takes as input taxonomic composition data (such as tables of Operational Taxonomic Unit (OTU) or Amplicon Sequence Variant (ASV) abundances), and links each component to its functional annotations through interrogation of the UniProt database. A functional profile of the gut microbiota is built from this basis. Both profiles, microbial and functional, are used to train Random Forest classifiers to discern unhealthy from control samples. An automatic variable selection is then performed on the basis of variable importance, and the method can be iterated until classification performances diminish. This process shows that the translation of the microbiota into functional profiles gives comparable, albeit slightly inferior performances when compared to microbial profiles. Through repetition, it also outputs a robust subset of discriminant variables. These selections were shown to be more reliable than those obtained by a state of the art method, and its contents were validated through a manual bibliographic research. The interconnections between selected OTUs and functional annotations were also analyzed, and revealed that important annotations emerge from the cumulated influence of non-selected OTUs.

## Introduction

The importance and perspectives opened by the human gut microbiota have been at the forefront of the discussion in the medical field in the past years, as a wide array of unsuspected impacts on host health have been derived from its composition. As such, an increasing amount of research aims to use it as a vector for understanding and potentially treating diseases [1], a large panel of which have thereby been correlated to the microbial composition of the gut microbiota [2]. Computational biology approaches have been employed due to the increasing availability of relevant data to predict the health state of individuals on the basis of their gut microbiota composition. Linear approaches based on differential expression, such as DESeq [3], have been vastly explored [4]. However, concerns were expressed about these approaches’ limitations when applied to highly complex data, such as the composition of the gut microbiota, and Machine-Learning (ML) approaches were proposed as a more adapted approach [5]. Studies in this vein [6–8] have found that relative abundances of taxonomic composition derived from gut microbiota sequencing made for an efficient predictor of host health status, with Random Forest (RF) [9] and Support Vector Machines (SVM) [10] emerging as the best adapted models.

In the medical community, discussion has emerged in the past years around the potential benefits of shifting from taxonomic analysis to understanding the functional aspects of the gut microbiota [11]. Many consider that host-microbiota interactions can only truly be understood on the functional level, and that comprehension of the microbiome on this scale is essential to envision strategies to improve host health via the gut microbiota. In this regard, taxonomic profiles such as Operational Taxonomic Unit (OTU) abundances provide insufficient information on their own, as OTUs can be functionally redundant and therefore have more relevance in regard to their cumulated influence on the community’s metabolome.

This article focuses on the question of the exploitation of functional gut microbiota profiles as a basis for health status classification, and how it compares to the already well-explored alternative based on microbial profiles. The comparison can be made on the basis of classification performance, but also on the subsequent potential for interpretability, with an enhanced focus on robustness and exhaustivity.

The shift to functional descriptions of the microbiota is possible through different approaches. One method is to take gene marker sequences or sequenced reads as input and effectuate a taxonomic assignation before matching it to functional annotations (FA), as done by tools such as PiCRUSt [12,13] or HUMAnN [14–16] to yield a measurement of the metabolic functions’ presence in the samples. Another approach is to directly explore datasets, from the basis of the microbial profiles, to gather information concerning each taxonomic affiliation’s functional profiling, as is implemented by the EsMeCaTa pipeline [17], which acts as a proxy for the interrogation of the UniProt database [18].

The exploitation of these functional profiling tools has allowed other studies to explore the potential of functional profiles as basis for the training of classifiers. Douglas et al. [19] converted sequenced samples, both issued from shotgun metagenomic sequencings (MGS) and 16S sequencing, from a cohort that includes individuals diagnosed with Crohn’s Diesease (CD) and healthy controls [20]. Jones et al. [21] converted MGS samples taken on pediatric CD patients, some of which have achieved sustained remission. Both studies then used these profiles as basis for training RF models to notably predict disease state or gravity, and response to treatment.

Performance-wise, Douglas et al. [19] compared the different profiles’ efficiencies when it comes to classification. In terms of accuracy first, for disease state classification, models trained on 16S information on OTUs, notably at the Genus and Phylum levels, were found to be significantly better than the others, while functional profiles were overall less performant, but still had significance in the case of 16S-derived orthologs and MGS-derived modules. Jones et al. [21] found that KEGG pathways could be reliable predictors when it came to treatment response when associated to complementary metadata. Douglas et al. [19] took a step further in comparing the profiles’ influences, as feature importances derived from combined random forest models confirmed that OTU abundances were overall more influent on disease state classification than annotations. Treatment response, however, had several pathways and orthologs in its top influent variables, showing that metabolic profiles also hold information.

In both of these studies, the shift to functional profiles is shown to decrease classification performance. However, they also illustrate the RF models’ potential for subsequent biological interpretation, notably exploiting the variable importance rankings of these models. Both studies included a discussion around the top features in terms of importance, and found them to be relevant with past litterature, both in terms of taxons and annotations. While these discussions served as validation for these works, they were not the main focus and only covered a limited number of variables: Douglas et al. [19] illustrate the top 30 features for each of their classifiers, and Jones et al. [21] discuss in detail the top 3 variables from each of their profiles. Little focus was put on the robstness of the results, with few repetitions of the experiments. These results do hint at the possibility to use RF classifiers as a means to highlight biologically relevant information from gut microbiota profiles however, and at the possibility to use a similar approach to establish a list of functions to rely on for further exploration of the question at the biological level.

The perspective of furthering these explorations on interpretability woul require an exhaustive approach. As such, a discussion mostly centered around top values selected based on an arbitrarily defined threshold, which introduces a human bias in the process of selecting the relevant information, would increase the risk of bypassing important variables. An automatic selection should therefore be prioritized. This exploration would also call for enhanced robustness in the approach, with more repetitions of the classifiers’ training process being a possible approach to this [22]. Finally, in exploring the gut microbiota through ML-driven approaches, the microbial and functional profiles are usually kept separate, and little discussion is made around the possibility to explore the interassociations and complementarity between microbial and metabolic signatures of the disease. This could be an interesting addition to the paradigm, as they would reveal the nature and prevalence of the functional cumulation of bacteria within samples.

In this article, we present a novel approach, implemented as an automated pipeline named SPARTA (Shifting Paradigms to Annotation Representation from Taxonomy to identify Archetypes), to explore and expand upon the capacities of the metabolic mechanisms of the microbiota in the context of a Machine Learning-driven approach. SPARTA retrieves the microbiota’s metabolic mechanisms, uses them as means to classify individuals with regard to their health status, and extracts significantly discriminating features from this process. This approach was tested on six different datasets previously used as reference for classification performance [7, 8]. A post-processing method is also implemented, to accentuate emphasis on genericity and robustness. This involves extracting an adaptative and robust shortlist of significantly discriminant features compiled from repetition of the method, which we backed with a manual bibliographic verification. Our pipeline also integrates and exploits the interconnections between bacteria and FAs, to facilitate biological interpretations of the outputs, as well as highlight the benefits of the latter profile.

## Results

### SPARTA overview: paired mechanistic analysis from relative microbial abundance profiles

SPARTA (see Fig 1) requires two inputs. The first is a table describing the microbial relative abundances (i.e: OTU abundance tables) for each microbiota sample within the dataset, from which functional profiles will be computed. The other is a vector file indicating the groups according to which each sample within the dataset should be classified, represented as green and red colors in Fig 1.

**Fig 1.**
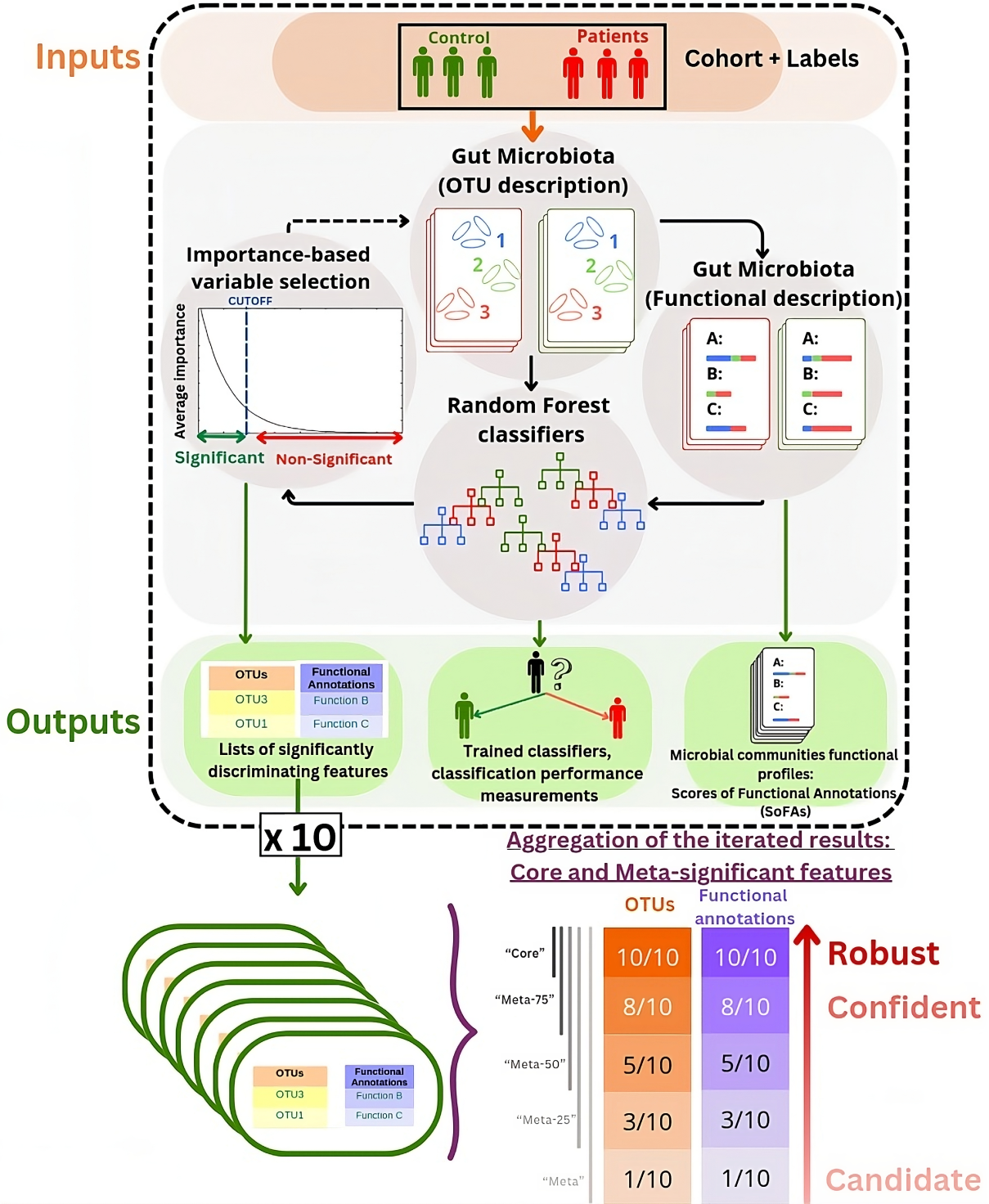
A schematic representation of SPARTA’s pipeline. From OTU tables and their associated labels as inputs, the pipeline produces functional descriptions of the microbiota samples via the EsMeCaTa pipeline. Both of these profiles are then used as basis for the training of Random Forest models to discern Control from Patient profiles. The average Gini importance scores of these variables over all trained forests is then used as basis for a selection of significantly discriminant variables, which can then be processed again iteratively, or passed as an output. For robustness, the process is repeated 10 times, leading to 10 different lists of significantly discriminant OTUs and FAs. These lists can be compiled into different categories, which group variables by level of robustness based on the frequency of their appearance in the significant lists. Thus, unanimous variables are considered to be “robust” discriminators, those agreed on by 75% or more of the classifiers are considered “confident”, and those that are selected at least once are considered “candidates”. Internally to the pipeline implementation, robust features are labeled “Core-significant”, and the others are labeled as “Meta-X significant”, X being the percentage of significant variable lists that include them.

SPARTA computes three major outputs. The first is a functional profile: by using the EsMeCaTa tool [17] to query the UniProt [18] database, we associate a representative proteome to each OTU from the original profiles, and link them to FAs (Gene Ontology (GO) terms [23], Enzyme Commission (EC) numbers [24]). The prevalence of each of the obtained annotations within the individual samples are then calculated as scores of FAs, as described in Materials and Methods.

The second consists of classification performances: SPARTA trains Random Forest (RF) [9] classifiers on the obtained functional profiles, and measures their performance in categorizing the samples.

Finally, SPARTA generates a list of features, both OTUs and FAs, which are identified as significantly discriminating between the given sample groups on the basis of an automatically calculated selection threshold applied to their average Gini importance scores [9] (see Materials and Methods). The associations between OTUs and annotations are also explicitated allowing each feature to be linked notably to its significant counterparts.

The pipeline can be iterated by selecting the information related to the shortlisted features in the initial dataset and repeating the steps. This process generates shortlists of significantly discriminating features that can be combined for a robust consensus. SPARTA is applied 10 times, each time with different test subsets, leading to some differences in variables considered significant. To address this, variables are categorized as follows: **(i)** “Robust” if unanimously deemed significant in all SPARTA runs (above the variable selection threshold). This category contains the variables that are most essential to the discernment of both patient profiles. **(ii)** “Confident” for the variables that were considered significant by at least 75% of the different runs (in our case, by 8 or more runs out of 10). This category contains variables that are likely to be important for profile discrimination, and could be a complement to the robust shortlist for interpretation. **(iii)** “Candidate” for variables shortlisted in at least one SPARTA run. These are variables that should not be fully excluded from consideration when it comes to interpretation, but that are unlikely to be influential. More generally, across all of these categories, the robustness of a selected variable can be evaluated in the light of the number of different SPARTA runs that list it as significantly discriminant.

Overall, OTUs and FAs are quantified on three different levels by SPART They are given: **(i)** A score based on their presence in each individual sample, in the form of a matrix containing, per sample, the relative abundances for OTUs (output ‘OTU table stripped.tsv’), or the scores for annotations (output ‘SoFA table.tsv’), **(ii)** A quantification of how discriminant they are between profiles of samples in the form of a vector of importance scores (outputs ‘OTU name of disease.csv’ and ‘scores name of disease.csv’ for OTUs and scores of FAs respectively), **(iii)** An indicator of their robustness as a discriminator, in the form of lists of variables affiliated to the “robust” and “candidate” categories (outputs [core/meta] name of disease run nb.cs

### Differential analysis of OTU and functional results

#### Experimentation

We applied SPARTA to six publicly available datasets, previously explored in articles covering approaches such as MetAML [8] or DeepMicro [7]. These datasets contain OTU abundance tables issued from sequenced microbiota samples from cohorts of healthy controls and individuals diagnosed with Cirrhosis (Cirrhosis dataset), Colorectal Cancer (Colorectal dataset), Obesity (Obesity), Type 2 Diabetes (T2D and WT2D datasets) or Inflammatory Bowel Disease (IBD dataset). For further details, see Materials and Methods. SPARTA was launched with the default parameters of 10 runs, 5 selection iterations per run, and 20 trained models per iteration. We analyzed the results in terms of classification performance and variable selection, as seen below. Further details are available in Supporting Table S1, and raw results are in Supporting File S1.

#### Machine Learning classification performances obtained from functional profiles are similar to those from OTU profiles

Fig 2 illustrates the classification performances of the Random Forests [9] trained by SPARTA to distinguish between patients and healthy individuals, per profile and dataset. The represented metric is the average AUC of the 20 Random Forests trained in this SPARTA iteration, as described in Materials and Methods. Only the optimal selection is represented here, defined as the selection level that maximizes the median of this metric over 10 repetitions. The number of iterated selections corresponding to this selection are given in the ‘Optimal Selection’ column. For each dataset, a Mann-Whitney U-test was conducted comparing the performances based on the OTU and functional profiles at respective optimal selection levels. For example: the Cirrhosis dataset’s functional (orange) and OTU (blue) profiles have been tested over 10 runs by SPARTA. These tests have allowed us to detect the level of variable selection that yields the best median classification scores for each profile, which were then chosen for this representation. In this case, as shown in the ‘Optimal selection’ column, both profiles hit peak performance after 1 iteration of variable selection. Each of the 10 runs of SPARTA yields an average classification performance score, corresponding to the plotted dots. The boxplots represent the associated distribution, and notably show that the Functional profile has a median averaged AUC of 0.92, against 0.95 for the OTU profile. The difference between both distributions was not found to be significant by a Mann-Whitney U-test, as shown by the absence of an asterisk symbol on this row.

**Fig 2.**
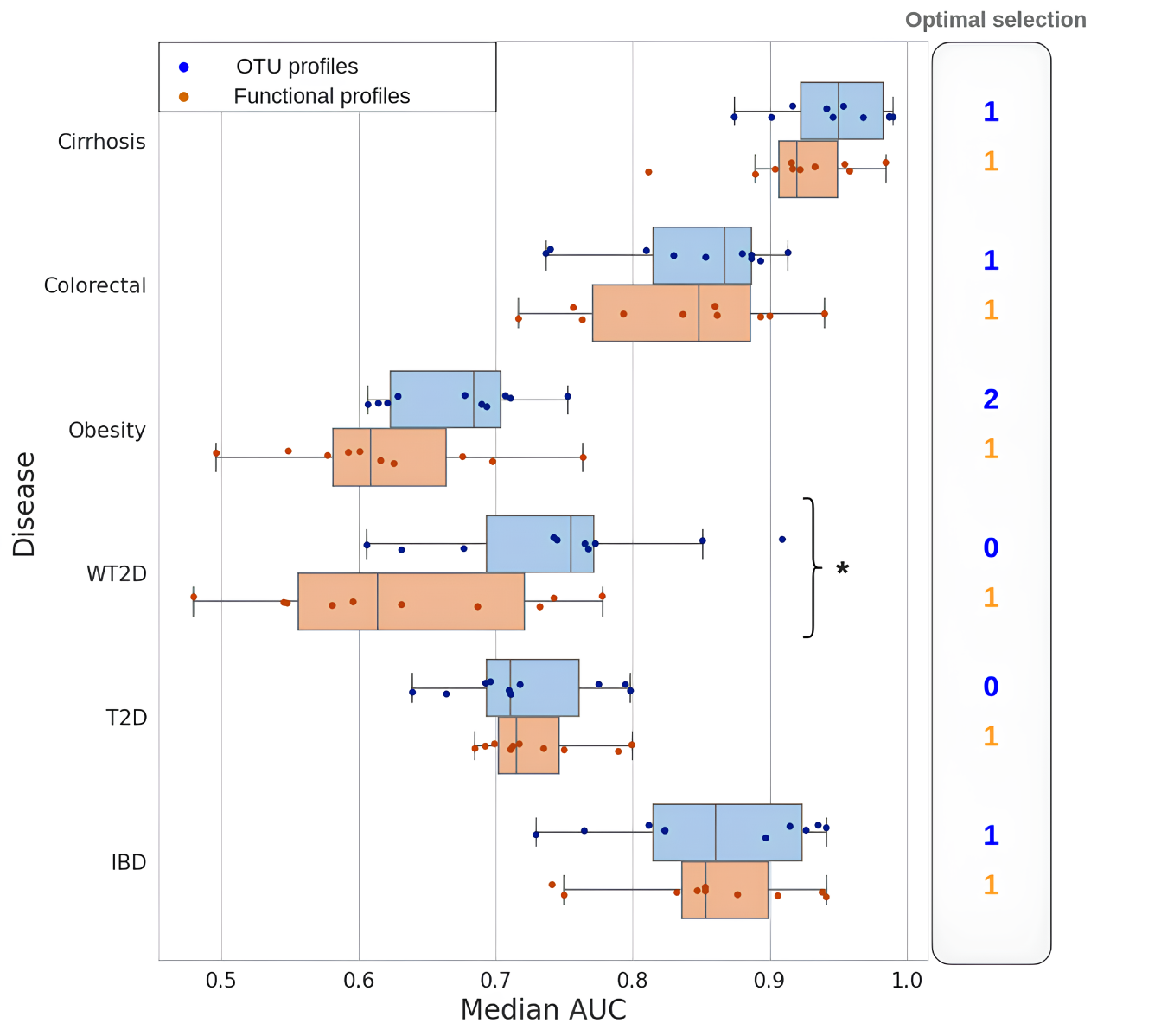
Classification performances of Random Forest models trained on OTU and functional profiles, and impact of the variable selection on performance. Median classification performances (AUC) for all types of profiles and each dataset, at the optimal level of selection over 10 full runs of the pipeline. Each of these runs involved a different randomly selected test set of individuals, which was used for both profiles. Performances and importance scores for each run were computed and averaged over 20 distinctly trained Random Forest models. The amount of selection iterations required to obtain the best average among these median AUCs are represented beside each plot. Profiles that have a significantly better classification performance than their counterpart for the same profile (based on a Mann-Whitney U-test) are signalled by a * symbol.

Overall, we can see that OTU profiles yield better median classification performances than their functional counterparts, with the T2D dataset being the only exception. However, the difference in performance between both profiles is only significant in the case of the WT2D dataset, showing that though converting our data to the functional level comes at the cost of some performance, both profiles perform comparably as basis for classification.

Of note is also the asymetrical benefit of variable selection. Functional profiles systematically benefit from a reduction of dimensionality, reaching peak performance after one selection, whereas variable selection only hinders classification performance in two of the six datasets for the OTU profiles.

These results are in line with the previous works of Douglas et al. [19] and Jones et al. [21] in that OTU profiles are here again shown to overall be a better basis for Random Forest prediction. However, the innovative potential of functional profiles resides more in their prospective contribution to a biological understanding of the diseases’ mechanisms than in their use for automatic classification. We will now focus on propositions to optimize the differential functional profiling of microbiotas in the context of a disease, as well as evaluate the added value of functional information in comparison to OTUs for understanding the underlying biological processes.

#### Robustness of SPARTA’s feature selection: comparative evaluation

The datasets used in the previous section contained on average 484 OTUs. Through EsMeCaTa’s [17] pipeline and its interrogation of UniProt [18], these OTUs were linked to a total average of 10,510 FAs per dataset, resulting in a 22-fold mean increase in the amount of information, as shown in Table 1. For example: in total, the sequenced samples of the Cirrhosis dataset covered 542 OTUs, which were associated by EsMeCaTa to a total of 10,434 FAs. Following SPARTA’s application, 69 of these OTUs and 970 of these annotations were included in the candidate sublists. Among these, 30 OTUs and 312 annotations were in the confident subset, and 21 OTUs and 209 annotations were in the robust subset.

**Table 1.** Application of the SPARTA selection process to identify signature OTUs and functions on 6 reference datasets. Total amount of features (OTUs and FAs) in the original dataset (“Initial Number” column) and in the robust, confident and candidate selections at the optimal SPARTA selection threshold (Calculated over 10 run of the pipeline).

| Dataset | Features | Initial number of OTUs | Predicted Functions | Robust subset | Confident subset | Candidate subset |
| --- | --- | --- | --- | --- | --- | --- |
| Cirrhosis | OTU | 542 | - | 21 | 30 | 69 |
|  | FAs | - | 10,434 | 209 | 312 | 970 |
| Colorectal | OTU | 503 | - | 24 | 46 | 116 |
|  | FAs | - | 10,635 | 53 | 190 | 2,225 |
| Obesity | OTU | 465 | - | 8 | 15 | 53 |
|  | FAs | - | 11,341 | 26 | 148 | 3,102 |
| WT2D | OTU | 381 | - | 381 | 381 | 381 |
|  | FAs | - | 10,180 | 6 | 59 | 3,082 |
| T2D | OTU | 572 | - | 572 | 572 | 572 |
|  | FAs | - | 10,275 | 128 | 279 | 1,485 |
| IBD | OTU | 443 | - | 26 | 37 | 102 |
|  | FAs | - | 10,196 | 64 | 158 | 1,631 |

To balance the afore described increase in information, SPARTA operates a selection of variables, as explained in Materials and Methods, aiming to correct issues that could stem from redundancy and dimensionality. This selection also generates one of the pipeline’s main outputs: a list of ranked features (either OTUs or FAs) based on their average Gini importance scores [9], and including an automatically computed cutoff that distinguishes significant and non-significant information.

The amount of information retained per SPARTA run for all functional datasets is illustrated by Fig 3A. The figure shows that the average amount of information to retain for optimal classification performance varies depending on the dataset. For example, retaining the top 500 annotations ranked by average Gini importance would give a selection similar to SPARTA on the IBD or Cirrhosis datasets, whereas the Colorectal dataset would require the top 700 annotations to match the selection. This shows that an adaptative method like SPARTA, which makes a decision concerning the quantity of information to be retained by the selection, has an advantage over a selection based on a fixed threshold, because it can adapt to the complexity of the problem at hand, which is shown here to be variable. SPARTA’s selection thresholds also do not match the more traditional thresholds, such as the top 30 features explored in Jones et al. [21], and can be used to get an estimate of the optimal amount of information to consider for discerning microbiota profiles.

**Fig 3.**
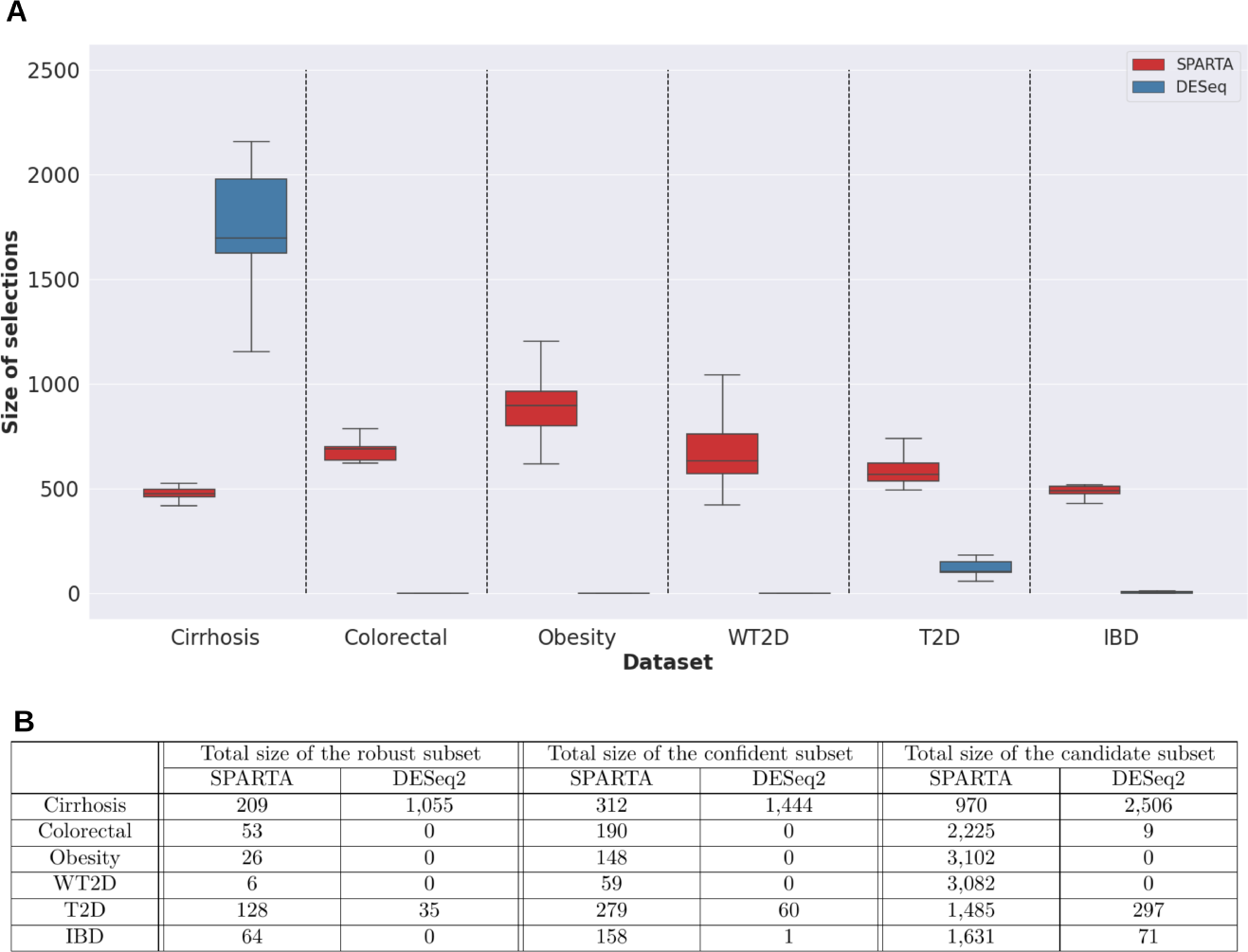
Comparative application of SPARTA and DESeq2 for variable selection. A: Amount of FAs selected by SPARTA and DESeq2, for all datasets. DESeq2 selections were effectuated with an adjusted p-value threshold of 0.01. Both selection methods were repeated 10 times, with a common test subset set aside each time. B: Sizes of the SPARTA and DESeq2 selections. DESeq2 was applied with an adjusted p-value threshold of 0.01. From left to right, the columns present, for SPARTA and DESeq2, the size of the robust, confident and candidate subsets issued by the concerned selection method iterated 10 time with identical test subsets.

Selections obtained from direct pairwise comparison of the profiles using the DESeq2 tool [3] were also explored. Variables were selected using a p-value threshold of 0.01, a classic threshold value exploited in several other studies that applied DESeq2 to metagenomic data [25–27]. Similarly to SPARTA, the selection process was iterated 10 times with variation induced from setting aside a subset of the samples, and variables were compiled into ‘robust’, ‘confident’ and ‘candidate’ categories depending on how often they were selected. Comparative results of this process are presented in Fig 3A and Fig 3B. For example, Fig 3A shows that, when applied 10 times to the Cirrhosis dataset, SPARTA selects a minimum of 417 annotations, and a maximum of 526, with a median of 474.5. In the same conditions, DESeq2 selects between 1,155 and 2,158 annotations, for a median of 1,696.5. These distributions are plotted, respectively, in red and blue. Fig 3B shows that with SPARTA’s selection, the Cirrhosis dataset’s outputs 209 robust annotations, 312 confidents, and 970 candidates, against a respective 1,055, 1,444 and 2,506 with DESeq2. With these parameters, DESeq2 is a much more stringent selector than SPARTA on all datasets aside from Cirrhosis. For the WT2D and Obesity datasets in particular, all selections are empty, leading to an empty candidate subset as described in Fig 3B. The IBD and Colorectal datasets also prove to be inadapted for this approach, yielding very small candidate and confident subsets, and empty robust subsets. Only the T2D and Cirrhosis datasets allow DESeq2 to yield a non-empty robust subset. SPARTA, on the other hand, consistently yields non-empty robust and confident selections, both of which are reasonably sized for interpretation when compared to the candidate subsets, being close to 50 times smaller in the case of the WT2D dataset’s confident and candidate subsets.

Focusing on the Cirrhosis and T2D datasets, which are the only datasets on which DESeq2 extracts a non-empty robust selection with an adjustedp-value threshold of 0.01, we have also explored the coherence between both approaches. Fig 4 illustrates the overlap between each method’s robust and candidate annotations.

**Fig 4.**
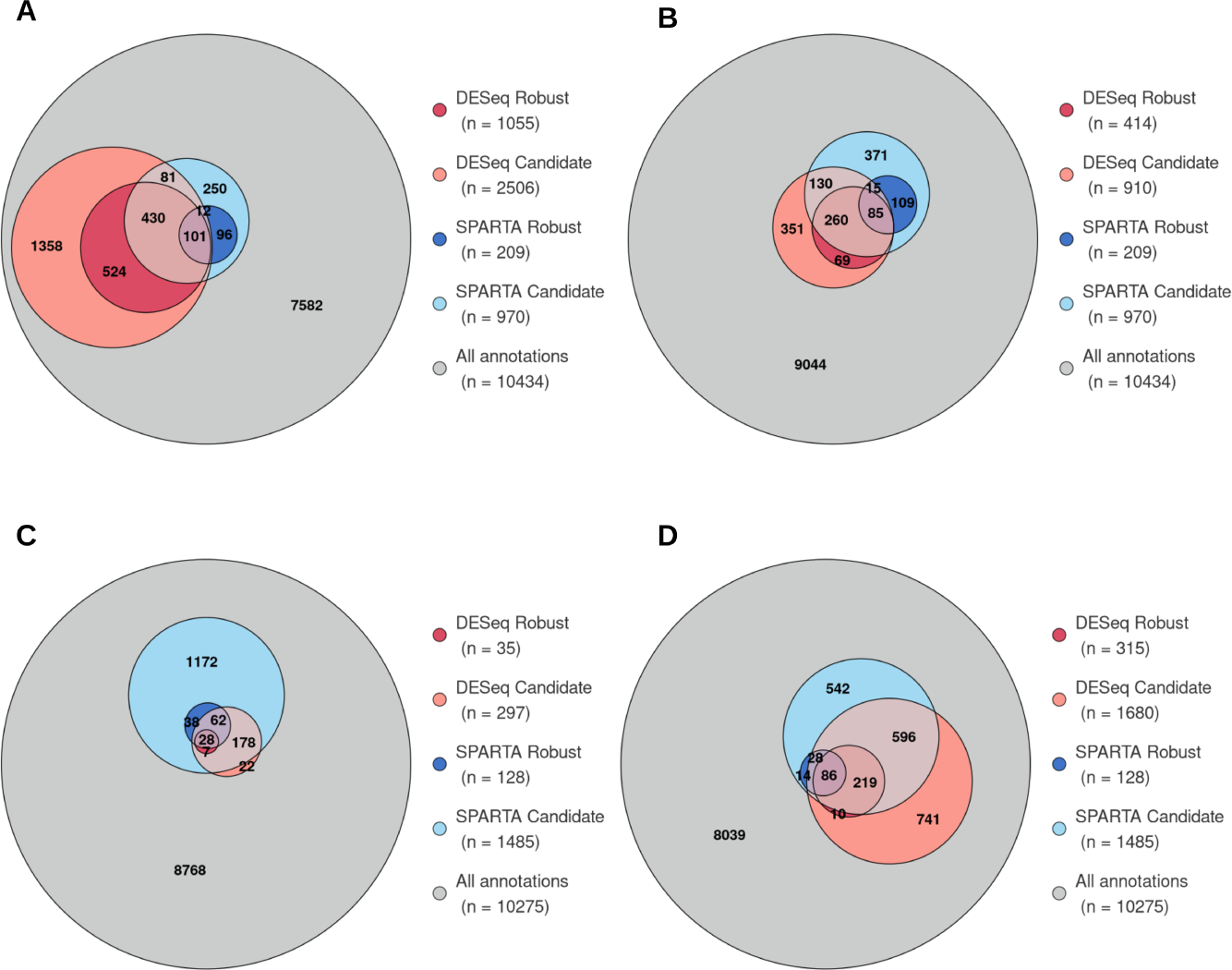
Robust and candidate lists overlap for SPARTA and DESeq2, obtained on the Cirrhosis (top) and T2D (bottom) datasets. The DESeq2 subsets illustrated on the left were obtained using the classic threshold of 0.01. Those on the right were obtained with a p-value threshold chosen to obtain comparably sized candidate sublists between SPARTA and DESeq2. Values indicate the number of annotations in each intersection, and do not represent the size of a category as a whole. A: Cirrhosis (adjusted p-value threshold = 0.01). B: Cirrhosis (adjusted p-value threshold = 1e-5). C: T2D (adjusted p-value threshold = 0.01) D: T2D (adjusted p-value threshold = 0.2). For representation of the confident subsets, see Supporting Fig S1.

T2D’s DESeq2 selection is smaller than SPARTA’s, englobing a total of 297 annotations in its candidate subset against 1,485 for SPARTA, as shown in Fig 3B. As shown by Fig 4C, only 22 of these annotations, representing 7% of the total size of the candidate subset, are not covered by SPARTA’s candidate selection. Also of note, the entirety of DESeq2’s robust subset is englobed by SPARTA’s selection, of which 80% by SPARTA’s robust selection. The Cirrhosis DESeq2 output, on the other hand, is bigger than its SPARTA counterpart, totaling 2,506 candidate annotations to SPARTA’s 970, as shown by Fig 4A. However, unlike in the previous case when the smaller selection here is quasi-englobed by the larger, SPARTA’s selection remains distinct from DESeq2’s, as 36% of the annotations in SPARTA’s total selection are not covered by DESeq2’s candidate subset. This is even more prevalent when zooming in on SPARTA’s robust subset, 46% of which is outside of DESeq2’s selection.

To put these results in perspective, there is no guarantee that a 0.01 p-value threshold yields an ‘optimal’ selection for both of these datasets when applying DESeq2. This choice of threshold is, however, a required external input for the method, that SPARTA does not need as it automates the choice of the selection’s size. As such, the chosen threshold could arguably be too restrictive for the T2D dataset, and too permissive for the Cirrhosis dataset. The latter is the more obvious outlier, firstly because we would expect a Random Forest model to select more variables than a method based on differential abundance like DESeq2, as it takes account of more criteria, and secondly because DESeq2’s total candidate selection accounts for a surprisingly high 24% of all annotations allocated to this dataset. This is further validated by the fact that, even though the Cirrhosis DESeq2 selection is much larger than SPARTA’s, a large proportion of the latter remains undetected by the former. Furthermore, the application of a more restrictive p-value threshold of 10^−5^, chosen to yield a total candidate selection for DESeq2 of comparable size to SPARTA’s, makes the Cirrhosis DESeq2 selection more similar to the SPARTA selection, as illustrated by

Fig 4B: 1,596 annotations are removed from the candidate selection due to the more stringent selection process, of which only 134 were in common with SPARTA’s selections, against 1,462 that were specific to DESeq2. A more restrictive DESeq2 selection therefore tends towards being incorporated in the SPARTA selection, supporting the hypothesis according to which SPARTA will select the information relevant to a DESeq2 selection of appropriate size, but will also take account of extra information that the former method cannot select, as it does not linearly differentiate the microbiota profiles. The same dynamic can be observed on the T2D dataset, on which a p-value threshold of 0.2, similarly obtained to generate a DESeq2 candidate selection of the same size as SPARTA’s, was applied, as illustrated by Fig 4D. This less restrictive threshold yields a DESeq2 selection that diverges from the SPARTA selection when compared to the first threshold, going from a 97% overlap with SPARTA’s candidates to 55%.

Overall, DESeq does not yield exploitable selections with a classic p-value threshold on four of our six datasets. The examination of the remaining two datasets allows us to illustrate how SPARTA and DESeq2 behave comparatively in different situations. In T2D’s situation, the DESeq2 selection is smaller and largely overlaps SPARTA, with DESeq2’s robust subset notably being entirely included in the SPARTA selection. For the Cirrhosis dataset, the SPARTA selection is the smallest of the two, however it remains distinct from what DESeq2 selects. A more restrictive p-value threshold makes the Cirrhosis DESeq2 selection tend towards inclusion in the SPARTA selection. This supports the hypothesis that SPARTA includes the top differentially expressed features of each dataset, but stops taking them in account at a selected, arguably optimized threshold, and takes into further account other features that differentiate the profiles in a non-linear manner.

### Exploiting biological knowledge from the paired robust functions and OTUs

For the following section, we will be relying on the robust outputs from the IBD dataset as an example. These results come from the pipeline’s first iteration, as it is the one that gives this dataset’s best classification performances on both OTU and functional profiles (see Fig 2). The IBD dataset was chosen as an average representative of our results, as it is an outlier in neither classification performance nor in the selection of variables by DESeq2.

#### Visualization of the robust shortlists

An important output of SPARTA is the shortlist of robust variables that are selected by the method, allowing for downstream interpretability. This comes in the form of tables of robustly significant annotations and OTUs, as previously described. The annotation shortlist for the IBD dataset is given in Table 2. It contains 64 FAs, alongside extra information that SPARTA helps associate to them. For example, annotation GO:0006520, corresponding to the amino acid metabolic process, is first in the table because it has the highest average Gini importance score over all 200 forests trained at this selection level, over 10 runs. It is on average 1.09 times as present in the diseased profiles as it is in the controls, the negative value of the ‘Ponderated average ratio’ meaning that the annotation is predominantly found in unhealthy samples. It is expressed by a total of 358 OTUs over all samples, of which 23 were found to be robust. The subsequent bibliographic analysis of this list graded its relevance to the disease as a 1, meaning that there is a known direct link between the annotation and IBD [28]. Detailed outputs are made available in Supporting File S2.

**Table 2.** Robust subset of annotations from the IBD dataset. Robust FAs of the IBD dataset, identified by their GO term or EC number, as well as their current name. Annotations are classified by decreasing average Gini importance score, over all 200 Random Forests trained at the optimal selection level (20 per run, 10 runs). Extra information include: the ratio between the average scores of the OTU in control and unhealthy profiles, ponderated by -1 if the OTU is most present in the unhealthy profiles, the amount of OTUs attached to each FA, and the amount of robust OTUs within them. Finally, the bibliographic category of each annotation, as defined in a subsequent section, is given.

| ID | Names | Average RF importance | Ponderated average ratio (Control/Unhealthy) | Number of linked OTUs |  | Bibliographic category |
| --- | --- | --- | --- | --- | --- | --- |
|  |  |  |  | Total | Robust |  |
| GO:0006520 | amino acid metabolic process | $4.96 \times 10^{-3}$ | -1.09 | 358 | 23 | 1 |
| 4.1.2.- | Aldehyde Lyases | $4.52 \times 10^{-3}$ | -3.21 | 28 | 2 | 2 |
| GO:0102545 | phosphatidyl phospholipase B activity | $3.49 \times 10^{-3}$ | -11.29 | 15 | 1 | 1 |
| GO:0008744 | L-xylulokinase activity | $2.86 \times 10^{-3}$ | -7.51 | 4 | 1 | 3 |
| GO:0004122 | cystathionine beta-synthase activity | $2.80 \times 10^{-3}$ | -7.40 | 8 | 1 | 1 |
| GO:0008788 | alpha, alpha-phosphotrehalase activity | $2.66 \times 10^{-3}$ | -4.05 | 19 | 1 | 3 |
| GO:0047419 | N-acetylgalactosamine-6-phosphate deacetylase activity | $2.56 \times 10^{-3}$ | -1.58 | 78 | 6 | 1 |
| GO:0001510 | RNA methylation | $2.42 \times 10^{-3}$ | 1.13 | 249 | 19 | 1 |
| GO:0016811 | hydrolase activity, acting on carbon-nitrogen (but not peptide) bonds, in linear amides | $2.41 \times 10^{-3}$ | -1.47 | 162 | 5 | 1 |
| GO:0015444 | P-type magnesium transporter activity | $2.36 \times 10^{-3}$ | -2.73 | 66 | 2 | 2 |
| GO:0008092 | cytoskeletal protein binding | $2.32 \times 10^{-3}$ | 3.23 | 3 | 1 | 1 |
| GO:0016832 | aldehyde-lyase activity | $2.27 \times 10^{-3}$ | -1.24 | 200 | 14 | 2 |
| GO:0032440 | 2-alkenal reductase [NAD(P)+] activity | $2.22 \times 10^{-3}$ | 3.29 | 5 | 1 | 3 |
| GO:0043130 | ubiquitin binding | $2.19 \times 10^{-3}$ | 3.18 | 6 | 1 | 1 |
| GO:0030798 | trans-aconitate 2-methyltransferase activity | $2.18 \times 10^{-3}$ | -1.79 | 18 | 1 | 1 |
| GO:0017065 | single-strand selective uracil DNA N-glycosylase activity | $2.16 \times 10^{-3}$ | 3.24 | 4 | 1 | 1 |
| GO:0042121 | alginate biosynthetic process | $2.04 \times 10^{-3}$ | 1.34 | 127 | 16 | 1 |
| GO:0045151 | acetoin biosynthetic process | $2.03 \times 10^{-3}$ | -1.89 | 48 | 1 | 3 |
| GO:1901135 | carbohydrate derivative metabolic process | $2.02 \times 10^{-3}$ | -1.20 | 271 | 17 | 1 |
| GO:0008707 | 4-phytase activity | $1.97 \times 10^{-3}$ | 3.24 | 1 | 1 | 3 |
| GO:0047605 | acetalactate decarboxylase activity | $1.95 \times 10^{-3}$ | -1.89 | 48 | 1 | 3 |
| 2.4.2.22 | xanthine phosphoribosyltransferase. | $1.92 \times 10^{-3}$ | -1.17 | 199 | 12 | 3 |
| GO:0008010 | kanamycin kinase activity | $1.91 \times 10^{-3}$ | -1.74 | 11 | 1 | 1 |
| GO:0005727 | extrachromosomal circular DNA | $1.87 \times 10^{-3}$ | -1.79 | 13 | 0 | 1 |
| GO:0003110 | xanthine phosphoribosyltransferase activity | $1.85 \times 10^{-3}$ | -1.79 | 201 | 12 | 3 |
| GO:0003951 | NAD+ kinase activity | $1.82 \times 10^{-3}$ | -1.08 | 337 | 19 | 1 |
| GO:0008610 | lipid biosynthetic process | $1.79 \times 10^{-3}$ | -1.98 | 87 | 1 | 1 |
| GO:0032265 | XMP salvage | $1.78 \times 10^{-3}$ | -1.17 | 199 | 12 | 2 |
| 4.1.1.5 | acetalactate decarboxylase. | $1.76 \times 10^{-3}$ | -1.89 | 86 | 1 | 3 |
| 2.7.1.51 | L-fuculokinase. | $1.75 \times 10^{-3}$ | 3.28 | 9 | 1 | 3 |
| 4.2.1.- | Hydro-Lyases | $1.74 \times 10^{-3}$ | -3.31 | 19 | 1 | 2 |
| GO:0008814 | citrate CoA-transferase activity | $1.67 \times 10^{-3}$ | -1.70 | 52 | 1 | 1 |
| GO:0016746 | acyltransferase activity | $1.66 \times 10^{-3}$ | 1.19 | 347 | 23 | 2 |
| GO:0046537 | 2,3-bisphosphoglycerate-independent phosphoglycerate mutase activity | $1.62 \times 10^{-3}$ | 1.15 | 185 | 20 | 3 |
| GO:0006741 | NADP biosynthetic process | $1.60 \times 10^{-3}$ | -1.08 | 329 | 19 | 1 |
| GO:0008760 | UDP-N-acetylglucosamine 1-carboxyvinyltransferase activity | $1.59 \times 10^{-3}$ | -1.11 | 361 | 20 | 3 |
| GO:0019677 | NAD catabolic process | $1.57 \times 10^{-3}$ | 1.63 | 32 | 3 | 1 |
| GO:0016297 | acyl-[acyl-carrier-protein] hydrolase activity | $1.54 \times 10^{-3}$ | 1.19 | 122 | 14 | 3 |
| GO:0006144 | purine nucleobase metabolic process | $1.54 \times 10^{-3}$ | -3.96 | 21 | 0 | 1 |
| 2.8.3.10 | citrate CoA-transferase. | $1.52 \times 10^{-3}$ | -1.70 | 52 | 1 | 1 |
| GO:0008453 | alanine-glyoxylate transaminase activity | $1.50 \times 10^{-3}$ | 3.22 | 5 | 1 | 3 |
| GO:0009346 | ATP-independent citrate lyase complex | $1.42 \times 10^{-3}$ | -1.72 | 52 | 1 | 2 |
| GO:0004135 | amylase-1,6-glucosidase activity | $1.37 \times 10^{-3}$ | 1.32 | 73 | 10 | 3 |
| GO:0003953 | NAD+ nucleosidase activity | $1.37 \times 10^{-3}$ | 1.66 | 29 | 3 | 1 |
| GO:0016977 | chitosanase activity | $1.36 \times 10^{-3}$ | 3.24 | 1 | 1 | 1 |
| 4.1.3.6 | citrate (pro-3S)-lyase. | $1.33 \times 10^{-3}$ | -1.72 | 52 | 1 | 1 |
| GO:0009409 | response to cold | $1.32 \times 10^{-3}$ | -1.48 | 35 | 0 | 1 |
| 1.8.4.10 | adenyllyl-sulfate reductase (thioredoxin). | $1.29 \times 10^{-3}$ | 2.43 | 21 | 2 | 1 |
| GO:0032049 | cardiolipin biosynthetic process | $1.27 \times 10^{-3}$ | 1.18 | 250 | 17 | 3 |
| GO:0047330 | polyphosphate-glucose phosphotransferase activity | $1.25 \times 10^{-3}$ | -6.70 | 5 | 1 | 1 |
| 3.5.3.6 | arginine deiminase. | $1.24 \times 10^{-3}$ | -1.78 | 58 | 2 | 1 |
| 4.1.1.31 | phosphoenolpyruvate carboxylase. | $1.21 \times 10^{-3}$ | -1.91 | 91 | 2 | 3 |
| GO:0008808 | cardiolipin synthase activity | $1.18 \times 10^{-3}$ | 1.17 | 239 | 17 | 3 |
| 1.1.1.22 | UDP-glucose 6-dehydrogenase. | $1.15 \times 10^{-3}$ | 1.29 | 142 | 14 | 1 |
| GO:0047356 | CDP-ribitol ribitolphosphotransferase activity | $1.15 \times 10^{-3}$ | -11.32 | 1 | 1 | 2 |
| GO:0097056 | selenocysteinyl-tRNA(Sec) biosynthetic process | $1.14 \times 10^{-3}$ | -1.42 | 214 | 6 | 1 |
| GO:0008964 | phosphoenolpyruvate carboxylase activity | $1.09 \times 10^{-3}$ | -1.91 | 91 | 2 | 2 |
| GO:0018800 | 5-oxopent-3-ene-1,2,5-tricarboxylate decarboxylase activity | $1.09 \times 10^{-3}$ | 3.38 | 8 | 1 | 3 |
| 2.7.1.23 | NAD(+) kinase. | $1.08 \times 10^{-3}$ | -1.07 | 325 | 19 | 1 |
| GO:0071702 | organic substance transport | $1.05 \times 10^{-3}$ | -1.41 | 103 | 6 | 3 |
| GO:0046110 | xanthine metabolic process | $1.03 \times 10^{-3}$ | -1.17 | 174 | 12 | 2 |
| 2.3.1.89 | tetrahydrolipicolinate N-acetyltransferase. | $1.03 \times 10^{-3}$ | -1.91 | 44 | 0 | 4 |
| GO:0005971 | ribonucleoside-diphosphate reductase complex | $1.00 \times 10^{-3}$ | -2.52 | 88 | 1 | 1 |
| 2.3.1.157 | glucosamine-1-phosphate N-acetyltransferase. | $8.18 \times 10^{-4}$ | -1.45 | 223 | 2 | 3 |

A similar selection of robustly discriminant OTUs is also available as an output of the pipeline, with the IBD output given as an example in Table 3. The same information as the previous table is available for each OTU, aside from the bibliographic categories. For instance, *Alistipes finegoldii*, identified in our process as OTU 73, similarly ranks first beacuse it has the highest Gini importance score on average over all trained Random Forests. Its differential expression shows that it is expressed on average 16 times as much in control profiles as it is in the unhealthy samples. As previously, we can establish which annotations are attached to each OTU, with *A.finegoldii* expressing a total 1,220 FAs, 16 of which are robustly significant. The details of these associations are available in Supporting File S3.

**Table 3.** Robust subset of OTUs from the IBD dataset. Robust OTUs of the IBD dataset, identified by their internal identifier, as well as their current name. OTUs are classified by decreasing average Gini importance score, over all 200 Random Forests trained at the optimal selection level (20 per run, 10 runs). Extra information include: the ratio between the average abundances of the OTU in control and unhealthy profiles, ponderated by -1 if the OTU is most present in the unhealthy profiles, the amount of FAs attached to each OTU, and the amount of robust annotations within them.

| ID | Names | Average RF importance | Ponderated average ratio (Control/Unhealthy) | Number of linked annotations |  |
| --- | --- | --- | --- | --- | --- |
|  |  |  |  | Total | Robust |
| OTU 73 | *Alistipes finegoldii* | $2.63 \times 10^{-2}$ | 16.40 | 1220 | 16 |
| OTU 224 | *Akkermansia muciniphila* | $1.99 \times 10^{-2}$ | 3.24 | 1452 | 24 |
| OTU 144 | *Lachnospiraceae bacterium 2 1 58FAA* | $1.87 \times 10^{-2}$ | -18.43 | 355 | 5 |
| OTU 12 | *Bifidobacterium bifidum* | $1.70 \times 10^{-2}$ | -11.32 | 1307 | 32 |
| OTU 171 | *Subdoligranulum unclassified* | $1.67 \times 10^{-2}$ | 1.56 | 627 | 6 |
| OTU 169 | *Ruminococcus lactaris* | $1.61 \times 10^{-2}$ | 3.04 | 431 | 3 |
| OTU 127 | *Eubacterium ventriosum* | $1.56 \times 10^{-2}$ | 2.64 | 1388 | 20 |
| OTU 134 | *Butyrivibrio unclassified* | $1.55 \times 10^{-2}$ | -1.65 | 916 | 9 |
| OTU 54 | *Odoribacter splanchnicus* | $1.52 \times 10^{-2}$ | 1.86 | 1595 | 20 |
| OTU 163 | *Ruminococcaceae bacterium D16* | $1.34 \times 10^{-2}$ | -4.30 | 1390 | 27 |
| OTU 156 | *Oscillibacter unclassified* | $1.29 \times 10^{-2}$ | -1.90 | 724 | 18 |
| OTU 78 | *Alistipes shahii* | $1.29 \times 10^{-2}$ | 1.65 | 935 | 8 |
| OTU 138 | *Coprococcus sp ART55 1* | $1.26 \times 10^{-2}$ | 2.27 | 701 | 8 |
| OTU 152 | *Roseburia hominis* | $1.23 \times 10^{-2}$ | 1.79 | 1500 | 20 |
| OTU 136 | *Coprococcus comes* | $1.09 \times 10^{-2}$ | -1.69 | 116 | 1 |
| OTU 75 | *Alistipes onderdonkii* | $1.07 \times 10^{-2}$ | 2.49 | 1391 | 16 |
| OTU 133 | *Butyrivibrio crossotus* | $1.07 \times 10^{-2}$ | 9789 | 1333 | 21 |
| OTU 53 | *Coprobacter fastidiosus* | $1.04 \times 10^{-2}$ | 6.07 | 1503 | 19 |
| OTU 162 | *Faecalibacterium prausnitzii* | $1.02 \times 10^{-2}$ | -1.57 | 1220 | 16 |
| OTU 154 | *Roseburia inulinivorans* | $9.62 \times 10^{-3}$ | 1.77 | 166 | 3 |
| OTU 30 | *Bacteroides cellulosilyticus* | $9.54 \times 10^{-3}$ | 2.28 | 345 | 3 |
| OTU 40 | *Bacteroides massiliensis* | $9.53 \times 10^{-3}$ | 1.54 | 1602 | 20 |
| OTU 74 | *Alistipes indistinctus* | $9.19 \times 10^{-3}$ | 1.27 | 1447 | 20 |
| OTU 20 | *Collinsella aerofaciens* | $9.13 \times 10^{-3}$ | -1.83 | 1349 | 22 |
| OTU 48 | *Bacteroides uniformis* | $8.49 \times 10^{-3}$ | 1.33 | 1809 | 17 |
| OTU 135 | *Coprococcus catus* | $7.83 \times 10^{-3}$ | 1.56 | 1496 | 25 |

#### Bibliographic exploration of the functional robust shortlist

Beyond the examples mentioned in this chapter, an in-depth bibliographic analysis of these outputs has been conducted for the IBD dataset, and is available in Supporting File S4.

The bibliographic examination was conducted on the integrality of the robust annotations from the IBD dataset, as well as samples of 20 annotations that were present in 50% of the significant sublists from SPARTA’s runs, and 20 non-candidate annotations. The methodology was to research the name of the annotation alongside the name of the disease on Google Scholar(https://scholar.google.com/). If none of the research results provided conclusive information linking this annotation to IBD, be it in a host model or in the microbiota, the chemical products of the annotation and eventual alternative names of the annotation were similarly tested, followed by related (parent or child) annotations, and finally the linked pathways listed in the BRENDA database [29]. From this exploration, the annotations were given a bibliographic relevance grade of 1 (most relevant to the disease) to 4 (least relevant to the disease) based on the following criteria:

Category 1: A direct link was established between the annotation, or a direct product metabolite, and IBD. This can come in the form of an explicitation of the metabolic mechanisms involved, or simply in the form of measured differential presence between unhealthy and control individuals. To note: conclusions derived from other ML-based approaches were not considered to be sufficient evidence, as they could suffer from biases similar to our own approach.

Category 2: A direct link was established between a similar metabolic function and the disease. Were considered as similar: proteins or enzymes from the same family as the one involved in the annotation (i.e: ATP-dependent and ATP-independent citrate lyases), and parent and child annotations, signaling notably that the annotation is indeed relevant, but at the wrong scale.

Category 3: An indirect correlation was established between the annotation and the disease. This can mean that the annotation was not directly linked to IBD, but that it is involved in a larger pathway or expressed in an OTU that has significance.

Category 4: No leads were found, or the annotation was proven to be irrelevant.

Among the robust annotations, several were found through bibliography to be relevant to the disease when expressed in the host organism as opposed to the microbiota. We considered both cases as a link found between the annotation and the disease, following the idea of permeability and interactions between the microbiota and its host [30].

When available, we also retrieved the group, namely unhealthy or control, most likely to express these annotations according to the bibliography. At the same time, SPARTA also retrieves the group that most expresses each of these robust FAs (see Materials and Methods). We confirmed these associations between FA and group with Deseq2 as well, for better robustness. We found that bibliography predictions and prevalence in the IBD dataset patients were in agreement in 58% cases. Functional annotations where disagreement exists between bilbliography and SPARTA/DeSeq might point towards a rescue of important functions in the host by the microbiota [31].

A complementary comparative analysis was conducted by the means of a Chi² contingency test with a 95% p-value threshold between the prevalences of each bibliographic categories in the robust selection and those of randomly selected non-candidate annotations (see Supporting Table S2 for details). This test established that the robust group significantly diverged from the non-candidate group. This significant difference is notably driven, as seen in Table S2, by a comparative increased proportion of Category 1, and decreased proportion of Category 4 annotations in the robust subset compared to the non-candidate selection. These results support the notion that SPARTA is a relevant selector of information.

#### Exploring the pairings between robust OTUs and annotations highlights their non-redundancy

Applying the pipepline to the IBD dataset (443 OTUs and 10,196 F.A.), the results show that annotations can be associated to 47.8 OTUs on average. One annotation is associated with the most OTUs (437 OTUs out of 443): GO:0016021, which is attached to the cellular membrane component, and is therefore expected to be extremely widespread. Unique associations account for 37.5% of all annotations, thus a majority of annotations are associated to more than one OTU. Overall, no function is perfectly ubiquitous, and the majority of functions are linked to several different OTUs. To quantify functional redundancy among OTUs, we used Jaccard proximity to measure the similarity of their functional associations. OTUs with a Jaccard proximity of 95% or more were considered functionally identical. Our analysis showed that 77.2% of the OTUs do not have such close neighbors, indicating that they maintain distinct functional profiles from each other, despite sharing many annotations with other OTUs. Detailed results are given in Supporting File S5.

The observed disparities between OTU and functional profilings prompt the question of whether these profiles equally provide valid descriptions of a subject’s microbiota. A potential drawback of the OTU scale is the cumulation effect, wherein individual OTUs may have little significance but contribute significantly to an essential metabolic process when grouped together. As a result, this collective impact might go unnoticed when focusing solely on individual OTUs. The dynamics in terms of specificity between annotations and OTUs are illustrated in Fig 5, which plots the amount of robust OTUs associated to each annotation as a function of the total amont of associated OTUs. For illustration purposes, the represented annotations were assigned into four profiles based on their number of associated OTUs. We labeled the top 10% as “Ubiquitous” (6 annotations, top right in Fig 5), the bottom 10% as ‘Specific’ (22 annotations, bottom left of Fig 5), and all others were labeled ‘In-Between’ (32 annotations). Finally, a fourth category was drawn up, independently of the previous criteria, containing 4 annotations that have no link to robust OTUs, which we labeled as ‘Cumulative’. This representation shows that important annotations have differing relationships to their OTU counterparts, and that an annotation’s importance can stem from the influence of several OTUs, as is notably illustrated by the ‘Cumulative’ class.

**Fig 5.**
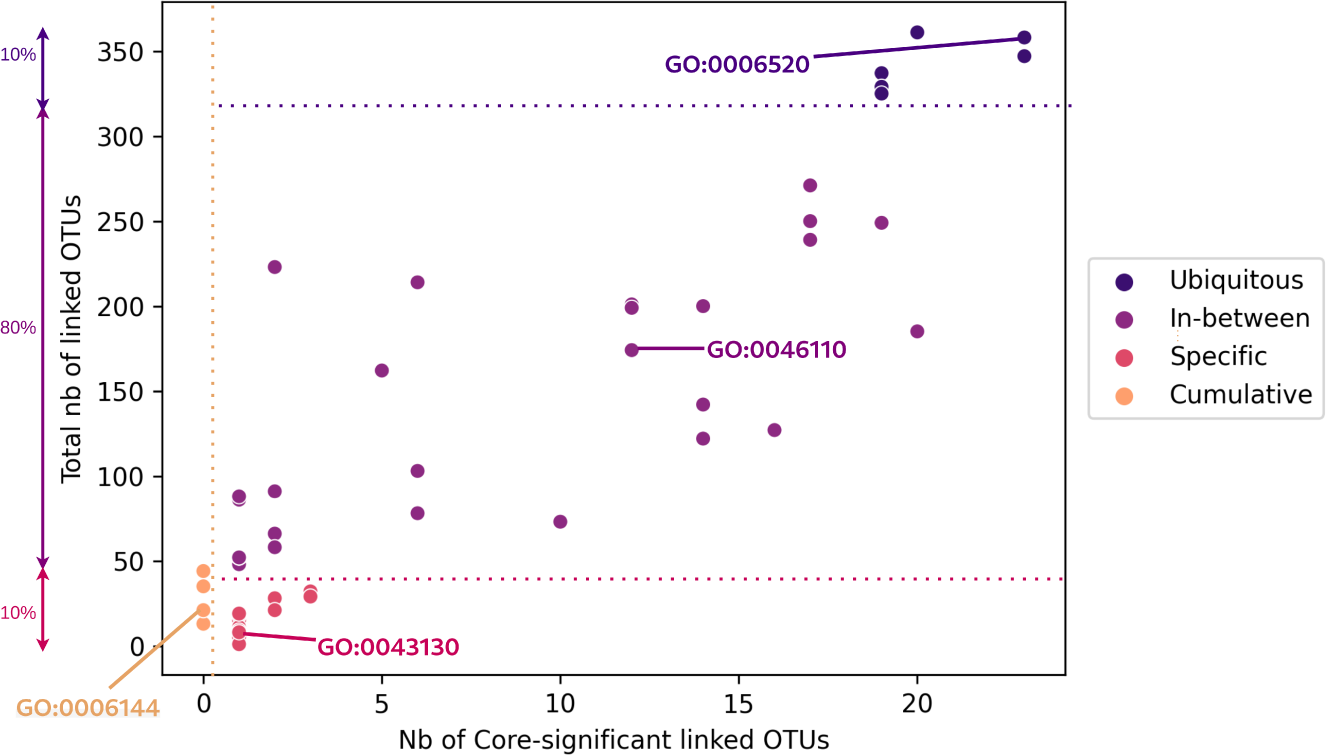
Number of OTUs associated to each robust annotation, as a function of the number of associated robust OTUs for the IBD dataset. Four groups of annotations are represented, three of which were determined based on the total amount of OTUs attached to the annotation: those within the top 10% of these values’ scale were labelled ‘Ubiquitous’, those in the bottom 10% were labelled ‘Specific’, and the others were labelled ‘In-between’. The final category corresponds to the robust significant annotations with no relationship to the robust significant OTUs (‘Cumulative’). The highlighted annotations are those used as illustrative examples in Fig 6.

A detailed illustration of annotations’ pairings with their OTU counterparts, as well as the strength of these links determined as described in Materials and Methods, is proposed in Fig 6. The represented annotations were picked each of the categories illustrated in Fig 5: GO:0006520 as representative of the ‘Ubiquitous’ class, GO:004610 for the ‘In-between’ class, GO:0043130 for the ‘Specific’ class, and GO:0006144 as a ‘Cumulative’ example.

**Fig 6.**
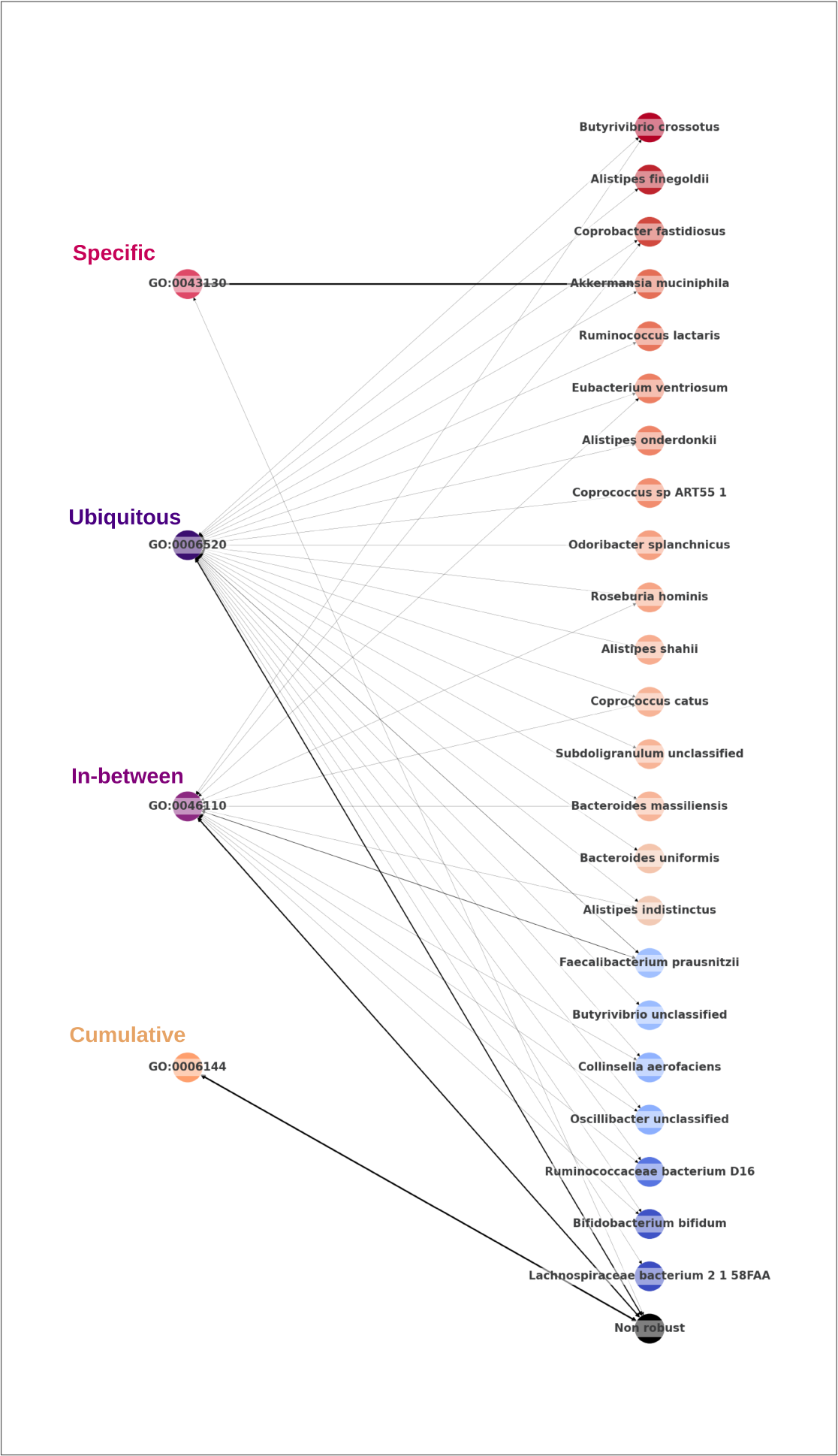
Associations between robust functions and the associated robust OTUs predicted by SPARTA, for the best iteration on the IBD dataset. Depicted annotations were selected to be representative examples of the different categories highlighted in Fig 5, and are presented with the same color scheme. OTUs are colored on the basis of their normalized average differential expression between Control (red) and Unhealthy (blue) profiles. The width of the connections is proportional to the importance of the association. The arrow between a given function and the generic ‘Non robust’ node represents the contribution of non-robust OTUs to the considered function.

From top to bottom on Fig 6, the first annotation (GO:0043130) is a case in which the feature’s significance appears to be due to a strong association to a single robust significant OTU, namely *Akkermansia muciniphilia*. This OTU has an established impact on IBD remission, and is researched as a potential probiotic treatment of the disease [32]. This is also in accordance with the annotation’s differential expression between profiles, as seen in Fig 6, where the annotation being colored in red shows that it is on average more expressed in control samples. This kind of relationship could either indicate that this ‘Specific’ annotation derives its importance in our predictions from its strong and specific attachment to an important OTU, or that its impact on the disease is an important factor to explain this OTU’s benefactory influence. GO:0043130 corresponds to ubiquitin binding, a mechanism which is known to regulate the inflammation process of intestines via different signalling pathways [33], and is categorized as a Category 1 annotation by our bibliographic research, showing that in the case of our example, the effects of the annotation and of its specifically associated robust OTU align. It should be noted that, as mentioned in our earlier discussion around our bibliographic work, the differential expression of a feature can be contradictory with its known effects, and should therefore be treated with caution. The second and third annotations (GO:0006520 and GO:0046110), respectively from the ‘Ubiquitous’ and ‘In-between’ groups, are very widespread among robust OTUs, without any particularly strong link to any of them. In cases such as these, meaning metabolic functionalities commonly expressed within OTUs, the issue of significance is shown to not be a purely binary question of expression or absence, as both annotations are consistently present in both profiles. Finally, the last annotation (GO:0006144) is exclusively linked to non robust OTUs. All such annotations are associated to several OTUs (13 minimum), meaning that their importance results from the cumulated influence of multiple, individually non-significant OTUs, that have a significant role when grouped functionally. The reverse associations, plotted in Supporting Figure S2, show that this form of cumulation is specific to FAs: the robustly significant OTU with the least associations to robust significant annotations, *Coprococcus comes*, is still shown to have a non-zero amount of correlations to robust annotations.

## Discussion

### From bacteria to functions

The first step of the SPARTA pipeline involves predicting annotations for the input taxonomic affiliations. To do so, we chose to rely on the EsMeCaTa pipeline, for the ease of its direct application to microbial profiles, as well as the presentation of its outputs which clearly explicitates the interassociations between taxonomic affiliations and FAs, making it more suitable for our subsequent analyses. Though our manipulations were made on OTU abundances, EsMeCaTa is also capable of processing Amplicon Sequence Variant (ASV) data.

It is however not the only tool available for the purpose of predicting functions, notably with the aforementioned PiCRUSt [12, 13] and HUMAnN [14, 15] pipelines, which are widely exploited. The cited works of Jones et al. [21] and Douglas et al. [19] notably rely on them.

These tools are more focused on calculating the presence of the functional annotations they attach to a sample, but put less emphasis than EsMeCaTa on keeping a trace of the interconnections between OTUs and functions, which makes them less suited for the latter parts of SPARTA’s approach. However, if we were to restrict our pipeline to its steps involving iterative Random Forest training and variable selection, a direct comparison to classification performances obtained from profiles built with PiCRUSt or HUMAnN could be envisaged for a more robust analysis of this aspect of the process.

### Comparative analysis of different approaches

#### Microbial and functional profiles

A central discussion point of this study is the pros and cons of exploiting the microbiome’s FA data as opposed to the explored OTU profiles for classification and interpretation.

In terms of classification performance, as shown in Fig 2 and discussed in Results, the OTU profiles remain a better overall predictor of disease state, though the difference is not significant in most cases. These results are in line with the findings of previous studies [19,21], and can be explained by the increase in the amount of features contained in the functional profiles. This is further supported by the fact that variable selection increases the functional profiles’ classification performances more consistently than the OTUs.

The main benefit of the functional profiles is that they are more in line with the current demands of the medical community [11] when it comes to the required precision level for biological interpretation. A potential caveat however would be the optimal amount of features retained by SPARTA, which greatly varies between both profiles as seen in Table 1, with the amount of annotations retained for optimal classification being consistently greater than the equivalent for OTUs. It seems intuitive that more metabolic functions would characterize unhealthy and control profiles when compared to OTUs, however the total amount of retained information in the case of FAs appears to be too extensive for biological interpretation to be practical. As such, we would recommend that interpretation of the FAs be limited on first approach to the robust and eventually confident subset outputs, which are in more manageable numbers, though these lists are unlikely to extensively cover all of the features relevant to the characterization of the disease.

#### SPARTA feature selection

SPARTA exploits the feature importance rankings that are inherent to the Random Forest method to perform a selection of variables. This selection step impacts classification performance, as discussed in Results, but is also important for the interpretability of the outputs, by highlighting the important elements within an otherwise overwhelmingly large list of features.

We also compared SPARTA’s selection to other approaches, as reported notably in Fig 3. By relying on an automatically computed cutoff threshold, our approach has proven to be more adaptative and robust than selections based on common fixed thresholds. The relevancy of exploiting Random Forests to perform selection as opposed to a more direct statistical comparison of unhealthy and control profiles was also highlighted when SPARTA’s selections are compared to those obtained with DESeq2, which measures differential expression. While it proved itself to be an efficient selector on datasets with clear distinguishing features, the latter tool did not detect any candidate features at realistic adjusted p-value thresholds when applied to a third of our test datasets, and did not have the internal coherence to generate a robust shortlist in another third. SPARTA on the other hand provided a robust subset for all datasets, showing it to be more consistent than DESeq2 when it comes to variable selection, especially in complex problems.

Random Forests are known to be capable of finding non-linear solutions to a problem [34], which explains the fact that a large amount of the information highlighted by SPARTA, including within the robust subset, remains undetected by DESeq even when the p-value threshold is unrealistically high, as shown by the results of Fig.4. As such, the content of SPARTA’s selection includes new information when compared to what can be extracted from linear comparisons.

#### Classification methods

SPARTA’s Machine Learing-based functionalities were based on Random Forest models, which are both a known performant classifier from the litterature [35], and a performant tool for interpretation as feature importances can be measured [9]. Other types of models have however proven to perform well in classifying individuals based on the gut microbiota, such as Support Vector Machines (SVM) [7, 8, 35]. While these classifiers are not as permissive to feature interpretability, a comparison of their classification performances on the data subsets highlighted by SPARTA with those obtained by the RFs could be a complement to our results.

### Post-processed outputs

#### Output interpretability

SPARTA’s end output is a shortlist of interconnected features, illustrated notably by the examples in Tables 2 and 3. The method puts emphasis on the selection’s robustness, as it is derived from the consensus of several repetitions, and adaptability, as the threshold for selection is based on an automatic calculation rather than a fixed rank selection. Its content also underwent bibliographic validation, in the case of the IBD dataset’s output. Though the list is likely not exhaustive, SPARTA’s selection was shown to be significantly enriched in bibliographically significant features. This supports SPARTA’s efficacy when it comes to highlighting factors that discriminate health profiles, though this should also be confirmed on the outputs obtained on other diseases.

The previously reported mismatches between the differential abundance-based profile attributions of SPARTA, which match those of DESeq2, and the conclusions of bibliographic research show that, in all probability, the underlying biological mechanisms involving these pathways are complex enough that a simple differential association is not sufficient to predict if an annotation is beneficial or detrimental to host health in the context of a given disease. A compensation mechanism could also be at play, as the gut microbiota is known to have the potential to compensate for metabolic functions that are lacking in the host [31]. A finer analysis of the Random Forest’s trained decision trees could give more appropriate insight into this issue.

It should also be noted that several annotations couldn’t be directly linked to IBD through bibliography (categories 3 and 4). These features deserve special attention, as they could be the result of a weakness of the method, or novel perspectives for research surrounding the disease.

#### Exploring links between OTUs and functions

Through Fig 5 and Fig 6, we established the reality of a cumulation effect, with OTUs that are less prevalent ending up having a detected influence on the microbiome’s metabolism through their combined contribution to a functional niche. This further supports the importance of exploiting microbiota information at the functional level rather than at the OTU level. Annotation GO:0006144, which corresponds to the purine metabolic process and is represented in orange in Fig 6, is a good illustration of this approach’s advantages. SPARTA’s outputs show that this annotation was not correlated to any robust OTU, and therefore would be difficult to derive from an OTU-based approach. Indeed, the bibliography shows that this annotation was linked to IBD through oriented research following a first mechanistic study [36], where our approach was capable of identifying it efficiently and without any pre-orientation.

### Applicability of the SPARTA pipeline and perspectives

Though it was tested on gut microbiota data, this method’s generic applicability can extend to other types of microbial communities. We focused on method robustness, presenting consolidated and exhaustive shortlists that showed agreement over 10 pipeline iterations without cherry-picking. These first results present a proof of concept for highlighting differentiating features in biological datasets through Machine Learning-based classification and variable selection, and establishing that integrating inter-associated taxons and functions for disease state classification with the gut microbiota enhances interpretability and exposes a functional cumulation effect. It also present opportunities for improvement. Integrating more specific external knowledge, such as individual metaclinical data, could enhance the interpretability of the questions for Machine Learning models. To further the comprehensiveness of our outputs and filter potential redundancies within annotations, we could explore leveraging Semantic Web information surrounding GO terms and EC numbers to aggregate or expand the existing information from UniProt. This could be the subject of future work.

## Materials and Methods

### The SPARTA pipeline: a Machine Learning-driven meth for paired analysis of taxonomic assignations and functional annotations

An implementation of SPARTA in Python, bash and R is available on github at https://github.com/baptisteruiz/SPARTA. The presented results were obtained from running in a Conda (version: 23.11.0) [37] environment that contains the EsMeCaTa pipeline (version at https://github.com/AuReMe/esmecata/tree/0.2.12) [17], as well as the following Python packages: pandas (version: 1.4.3) [38], numpy (version: 1.21.2) [39], scikitlearn (version: 1.1.1) [40], matplotlib (version: 3.5.1) [41], joblib (version: 1.1.0) [42], seaborn (version: 0.12.2) [43], progress (version: 1.6) [44], goatools (version: 1.2.3) [45], Biopython (version: 1.79) [46], requests (version: [47], kneebow (version: 1.0.1) [48], tensorflow (version: 2.4.1) [49] and keras (version: 2.4.3) [50].

The entirety of the pipeline can be launched with the ./sparta_main.sh-d name_of_input_files command, with the following optional arguments:

- -t: data treatment (can be: ‘tf igm’ or ‘scaling’, default: no treatment)
- -s: transform abundances to relative (can be: ‘relative’, default: no transformation)
- -i: number of iterations of the method (default: 1 iteration)
- -f: amount of trained classifiers per iteration of the command (default: 20 forests)
- -r: amount of pipeline runs (default: 1 run)
- -e: launch EsMeCaTa within the pipeline (default: True)

The pipeline is executed in four steps. The first is a run of the EsMeCaTa pipeline [17], preceded by formatting steps for the creation and formalisation of EsMeCaTa’s input from the given data. This step exploits the pipeline as described in a following section.

The second step is the calculation of the scores of the FAs obtained this way, following the method described further down and using the list of associations between taxons and annotations, as well as the original microbial abundances. This step can also involve data treatment, in accordance with the arguments parsed in the command line. If the ‘-s’ argument is given the ‘relative’ value, the taxonomic abundance profile will be forcefully converted to a relative abundance profile, with each value being recalculated as a percentage of the sample’s total, before the functional profile is calculated. Once we have the functional profile, its values can also be scaled, either using sklearn’s [40] ‘StandardScaler’ in the case where the ‘-t’ argument is assigned ‘scaling’, or TF-IGM, as described further in the Methods section, if the argument is given the ‘tf igm’ value. At this point, samples to be separated as a test dataset are also chosen, as a random sample that respects the original dataset’s label distribution.

The third step involves the training of 20 successive Random Forest classifiers to sort individuals according to their associated labels (i.e: ‘sick’ or ‘control’), based on the relative abundance profiles of their microbiota or on their calculated mechanistic representation. Before any training, a subsample of 20% the size of the full dataset is set aside as a test set. During training, the remaining data is randomly split into a training set and a validation set, with a respective 80% / 20% distribution. In order to account for the disparity in representation between the unhealthy and control individuals within the datasets, both classes were given weights proportional to their frequency, as implemented by scikit-learn’s ‘balanced’ class weight parametre [40]. The training involves a Grid Search, as implemented by scikit-learn [40], to optimize the estimator’s parametres in terms of the number of estimators per forest, the number of leaves per estimator, and the amount of information to which each tree has access. The split quality criterion is measured via the Gini Impurity metric. This step exports a list of each trained forest’s features’ Gini importances [9], as well as their classification performances (see Results) on the validation and test datasets. The best performing model on the test set is also exported.

The final step involves averaging all features’ importance scores over 20 training iterations, and selecting which ones are ‘Significant’ through a cutoff at the significance threshold, calculated as described further on. The list of all features listed by decreasing importance, with the position of the cutoff threshold, is then given as output.

If the ‘-i’ argument is given a value higher than 1, a subset of the original microbial and functional profile files will be created containing only the ‘Significant’ data. In the case where a data treatment method was given as input (‘scaling’ or ‘tf igm’), this step will be re-applied to the subset. After this, step 3 and onwards will be repeated for as many times as demanded through the ‘-i’ argument.

The entire process will be repeated, with the same paramaters, for as many times as dictated by the ‘-r’ argument. Each of these runs will have a new subset of test individuals. Once all requested runs have been completed, the shortlists obtained by all runs for each iteration are combined to categorize taxons and annotations as ‘Robust’ (outlined as significant by all predictors for a given iteration), ‘Confident’ (outlined as significant by at least 75 % of predictors for a given iteration) or ‘Candidate’ (outlined as significant by at least one predictor for a given iteration).

### Shifting representations, from microbial to functional profiles

The first step of SPARTA’s process is to transition from a representation of the microbiota on the scale of taxonomic affiliations to that of biological functions, by calculating the abundances of the FAs linked to the input’s taxons. In parallel, we are aiming to conserve the information linking together taxons and annotations, to expand upon this information later on. We also used the normalization of the annotation scores to introduce an *a priori* bias to boost the profiles of the best differentiating variables, in anticipation of the following classifying tasks.

#### Associating functional annotations to taxonomic affiliations: the EsMeCaTa pipeline

The EsMeCaTa pipeline follows three steps. The first step, ‘proteomes’, takes as input a tabular that associates a given name for all the studied bacteria to their exact taxonomy. From this, EsMeCaTa interrogates the UniProt database for proteomes associated with the taxon in question. If none can be found, the step is re-iterated with the superior taxonomic rank, until at least one proteome can be associated with the unit. In the event that more than 99 proteomes are associated with an taxon, a random selection of around 99 proteomes will be made, with respect to the taxonomic diveristy of the initial proteomes set. The selected proteomes are then downloaded from UniProt.

The second step, ‘clustering’, selects protein clusters that are representative of the taxonomic unit within the downloaded proteomes. To do so, the MMseqs2 tools [51] is used to create clusters of similar proteins from the proteomes. If a protein cluster contains similar proteins from 95% of the proteomes attributed to the taxonomic unit, it will be retained as part of its meta-proteome.

The final step, ‘annotation’, fetches the FAs (GO terms and EC numbers) of the retained protein clusters by interrogating the UniProt databases The final output is an ensemble of tabulars, one per taxonomic affiliation in the input, that contains all of the protein clusters kept in the taxon’s meta-proteome and their FAs.

#### Calculating a functional representation of the patient’s microbiota from taxon-annotation pairings

In order to compute a representation of the gut microbiota on the scale of the FAs, mixing information concerning its specific composition with the associated metabolic mechanisms, we give each annotation (F) a score, labeled as a Score of FA (SoFA), within a subject sample (i), similarly to [12], according to the following formula:

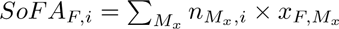

where *n_M_x_,i_* is the abundance value of taxon x within sample i, and *x_F,M_x__* is the number of proteins within taxon x’s proteome that are linked to the function F.

#### Normalizing and scaling data based on expected relevance with TF-IGM

The TF-IGM method [52] is used to normalize the results presented in this article. It was originally exploited in Natural Language Processing, as a method to highlight terms in a corpus of texts that are significantly present within a text while penalising those that are too widespread. The formula had to be re-adapted to fit our data and circumstances, and in our pipeline it is calculated based on the following two components:

- TF (Term Frequency): equivalent to the frequence of an annotation within the totality of a sample i:

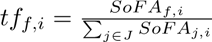 where *SoFA_f,i_*is annotation f’s score within sample i, and J is the ensemble of the annotations recorded within sample i.
- IGM (Inverse Gravity Moment): for each annotation f, the calculated values for *tf_f,i_* are ranked in decreasing order and noted as *T* (*f*)_1_,…,*T* (*f*)*_n_*, so that *T* (*f*)_1_ *> T* (*f*)_2_ *> … > T* (*f*)*_n_*, n being the total number of samples. We then have:

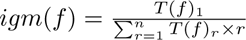 where r is the rank of the T(f) score in the previously defined order. The total TF-IGM score of an annotation f within a sample i will then be:

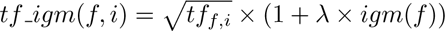 where *λ* is a value between 5 and 9. As per Chin et al.’s [52] recommendation, its value was set to 7 by default.

### SPARTA characterizes sample profiles with variables highlighted on the basis of a non-linear approach

Having established two types of profiling for microbiotas, we then explore their potential in differentiating classes, such as individuals based on their health status. To account for the complex interdependencies of biological pathways in impacting host health, we relied on Machine Learning classifiers rather than linear statistical approaches to establish the relevance of variables when it comes to distinguishing unhealthy individuals from controls. A method for robust selection is also proposed here, with an automated shortlisting of variables based on their importance, and a compilation of results accounting for consensus across multiple iterations of the method.

#### Training of Random Forest models

A Random Forest [9] classifier is trained to sort individuals in two classes (here, patients or controls), based on the relative abundance profiles of their microbiota or on their calculated mechanistic representation. Before any training, a subsample of 20% the size of the full dataset is set aside as a test set. During training, the remaining data is randomly split into a training set and a validation set, with a respective 80% / 20% distribution. In order to account for the disparity in representation between the unhealthy and control individuals within the datasets, both classes were given weights proportional to their frequence, as implemented by scikit-learn’s ‘balanced’ class weight parameter [40]. When measuring the performance of our classification algorithms, the metrics used were the Area Under the Receiver Operating Characteristic Curve (AUC) [53] averaged over 20 training iterations.

#### Extracting significant information from trained classifiers

Following the classifier’s training, the resulting feature importances are extracted. These importances are based on the Gini Importance metric, calculating the mean accumulation of the impurity decrease within each tree, as implemented in the Scikit-learn Python library [40]. If multiple iterations of the classifier’s training are made, the feature importances are averaged over all iterations. Features are then ranked based on this metric in decreasing order.

Once ordered, we aim to distinguish a separation between the features that were essential to the clasifier’s functionality, and those with a lesser impact. We place this threshold at the inflection point of the curve representing the decreasing importance scores, determined via an implementation of the Kneebow method [48], with all features above this point being labeled as “Significant”, and those below as “Non Significant”.

This process is iterated 5 times by default by SPARTA, and the optimal level of selection that is retained is the one that yields the highest median AUC during the classification process over 10 iterations of the pipeline.

#### Repetition of the SPARTA pipeline

In order to obtain robust results, the process of selecting a test subset, training classifiers and extracting significant features for a set amount of iterations, was repeated over 10 runs in our manipulations. Variations in the training conditions, with different test subsets selected for each run, result in 10 different shortlists of significant features per iteration. We label as ‘Robust’ the features that constitute the intersection of these shortlists, as ‘Confident’ those that are present in 75% or more of them, and as ‘Candidate’ those that are present in at least one of them. The amount of times a variable is labeled as significant by the optimal level of selection is an indicator of how reliable it is for the distinction of the differentiated profiles.

### SPARTA explicitates and quantifies the pairings between significant variables

Beyond significant shortlists, SPARTA also aims to illustrate the links between bacteria and FAs. EsMeCaTa’s outputs list all of the annotations all of the annotations estimated to be expressed by each taxonomic affiliation in the database, as discussed in a previous Methods section. From this, we can establish the reciprocal association, linking all annotations to the taxons that express them. In order to quantify the reciprocal impact of a taxon on an annotation’s score, we can calculate the following score:

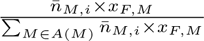

where *x_F,M_x__* is the number of proteins within taxon x’s proteome that are linked to the function F, 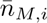 is the average of the abundances of a taxonomic affiliation within a dataset and A(M) is the ensemble of all taxons associated with the annotation.

### Assigning a feature to a profile

SPARTA also involves associating taxonomic affiliations and functional annotations to either the unhealthy or control categories. To do so, the profiles (relative abundances for taxons, scores of FAs for annotations) of all individuals within a same category were averaged, and the features were associated to the profile where they were most prevalent on average.

### Datasets

SPARTA was tested and benchmarked using publicly available specieslevel abundance profile datasets from the MetAML repository [8] and postprocessed for DeepMicro [7], concerning subjects diagnosed with a variety of diseases: Cirrhosis [54], Colorectal Cancer [55], Inflammatory Bowel Disease (IBD) [56], Obesity [57], and Type 2 Diabetes (T2D) on a Chinese [58] and an European [59] cohort. Each subject in these datasets had their gut microbiota sampled and sequenced with whole-genome shotgun and Illumina paired-end sequencing. The results were processed as per the standard procedure described by the Human Microbiome Project [60], then converted to species-level relative abundance profiles via the MetaPhlAn2 tool [61] with default parameters. Sub-species level features were then filtered using the MetAML tool [8].

Each cohort includes a portion of healthy control individuals, in addition to those who suffer from the disease in question. The proportions of each group in our cohorts are detailed in Table 4.

**Table 4.** Distribution of samples within the datasets of reference.

| Disease | Dataset | Total samples | Control samples | Patient samples |
| --- | --- | --- | --- | --- |
| Liver Cirrhosis | Cirrhosis | 232 | 114 | 118 |
| Colorectal Cancer | Colorectal | 121 | 73 | 48 |
| Inflammatory Bowel Disease | IBD | 110 | 85 | 25 |
| Obesity | Obesity | 253 | 89 | 164 |
| Type 2 Diabetes | WT2D (European Women Cohort) | 96 | 43 | 53 |
|  | T2D (Chinese Cohort) | 344 | 174 | 170 |

## Supplemental material

**S1 File. Detailed classification performances per dataset, SPART run and selection iteration.** The first sheet contains the detailed information as plotted in Fig 2: the average AUC scores, per run, for the overall best iteration level, for each dataset (OTUs and annotations). The median of the average values, and p-values of the Mann-Whitney U-test comparisons between the OTU and FA average scores per disease are also given. P-values under the 0.05 threshold are considered significant, and are highlighted with a *. Other sheets contain the details of each individual RF trained per run and iteration for each dataset (read: [dataset] R [run number] It [iteration number]). The information given per sheet is: for each of the 20 RFs trained this iteration and run, the optimal parameters found through GridSearch, the optimal threshold for probability prediction, and the AUCs on the training, validation, and test subsets.

**S2 File. Detailed robust and candidate functional annotation shortlists per dataset.** Each annotation is identified by its GO term or EC number, as well as its name. The complementary information given includes: the annotation’s average Gini importance over all RF models (‘Average importance’), the list of all OTUs associated to the annotation (‘Linked OTUs’) and the sublist of robustly significant OTUs within them (‘Significant linked OTUs’), and the profile it is associated to (‘Family’) supported by the average scores of the annotation in the patient and control samples. Outside of the robust shortlists, the number of SPARTA iterations that deem the annotation significant is also given (‘Count’).

**S3 File. Detailed robust and candidate OTU shortlists per datase** Each OTU is identified by its internal identifier (‘ID’), as well as its full taxonomy. The complementary information given includes: the OTU’s average Gini importance over all RF models (‘Average importance’), the list of all annotations associated to the OTU (‘Linked Reactions’) and the sublist of robustly significant annotations within them (‘Significant linked Reactions’ and the profile it is associated to (‘Family’) supported by the average abundances of the OTU in the patient and control samples. Outside of the robust shortlists, the number of SPARTA iterations that deem the OTU significant is also given (‘Count’).

**S4 File. Bibliographic exploration of the IBD dataset’s shortlists.** The detailed conclusions of the bibliographic research on IBD’s whole robust output, as well as random selections of 20 annotations that were non-candidates, and significant in 50% of SPARTA’s runs. Bibliographic categories are as presented in Results. The categorizations are justified by quoted sources.

**S5 File. Details of the pairwise Jaccard distance measurements between OTUs based on their associated annotations.** Pairwise Jaccard distances between OTUs, calculated on the basis of their functional profiles as detailed in the ‘Detail of OTU to annot links’sheet. The final column, ‘Sum of close neighbors’, counts the number of OTUs with a distance of 0.05 or less from the one concerned. A value of 1 in this column means that the OTU in question only has itself for a neighbor.

**S1 Table. Evolution of the average median AUC scores per dataset at increasing levels of variable selection, for taxonomic (OTU) and functional (SoFA) profiles.** The ‘No Selection’ column for each profile serves as a baseline for performance. The top performing selection levels are highlighted in bold.

**S2 Table. Counts of the different bibliographic categories per researched selection, and p-values of a Chi² contigency test compared to the robust subset.**

**S1 Fig. Robust, confident and candidate shortlist overlaps for SPARTA and DESeq2 selections on the T2D and Cirrhosis dataset** For readability, these representations were not scaled to the total number of annotations, as in Fig 4. A: Cirrhosis (adjusted p-value threshold = 0.01). B: Cirrhosis (adjusted p-value threshold = 1e-5). C: T2D (adjusted p-value threshold = 0.01) D: T2D (adjusted p-value threshold = 0.2).

**S2 Fig. Associations between robust OTUs and the associated robust functions predicted by SPARTA, for the best iteration on the IBD dataset.** Similarly to Fig 6, the colorscale for the OTUs is based on their differential expression between control and unhealthy profiles, and arrow length is proportional to the strength of the OTU’s connection to the annotation. Relationships to non-robust annotations were not represented here for reasons pertaining to readability of the figure. Represented OTUs were chosen to showcase control and healthy representatives with high and low numbers of connections to robust annotations.

## Supporting information

Supplemental Archive Description

## Acknowledgments

We would like to thank the GenOuest platform, which provided the computing resources used for obtaining the presented results. We would also like to thank Pauline Girard, Jeanne Got and Olivier Dameron for discussion and comments on the development of the method. First author was supported by a joint INRIA/INRAE PhD fellowship.

## Author Contributions

**Conceptualization:** Baptiste Ruiz, Anne Siegel, Yann Le Cunff

**Data Curation:** Baptiste Ruiz, Arnaud Belcour

**Formal Analysis:** Baptiste Ruiz, Anne Siegel, Yann Le Cunff

**Funding Acquisition:** Sylvie Buffet-Bataillon, Isabelle Le Hüerou-Luron, Anne Siegel, Yann Le Cunff

**Investigation:** Baptiste Ruiz

**Methodology:** Baptiste Ruiz, Arnaud Belcour, Samuel Blanquart, Sylvie Buffet-Bataillon, Isabelle Le Hüerou-Luron, Anne Siegel, Yann Le Cunff

**Software:** Baptiste Ruiz, Arnaud Belcour

**Supervision:** Anne Siegel, Yann Le Cunff

**Validation:** Baptiste Ruiz, Yann Le Cunff

**Visualization:** Baptiste Ruiz, Sylvie Buffet-Bataillon, Isabelle Le Hüerou-Luron, Anne Siegel, Yann Le Cunff

**Writing – Original Draft Preparation:** Baptiste Ruiz, Anne Siegel, Yann Le Cunff

**Writing – Review & Editing:** Baptiste Ruiz, Arnaud Belcour, Samuel Blanquart, Sylvie Buffet-Bataillon, Isabelle Le Hüerou-Luron, Anne Siegel, Yann Le Cunff

