## Supplemental Archive Description for "SPARTA: Interpretable functional classification of microbiomes and detection of hidden cumulative effects"

### Contents of the Supplementary Archive

March 1, 2024

#### 1 Accessing the Supplementary Archive

This archive is hosted on , at the following DOI: `10.5281/zenodo.10728698`. It contains material for reproduction of the article's figures, as well as detailed outputs from the SPARTA runs presented in the article, all within the 'SPARTA_reproduction_outputs.tar.gz' archive, alongside this supplementary document.

#### 2 Datasets

All datasets are made available in the accompanying S1 folder, with both the OTU tables and the label files derived from them. The OTU tables were originally available in the DeepMicro repository (https://github.com/minoh0201/DeepMicro). The original sources of the data are listed in Table S1.

| Disease | Dataset | Reference | Original reads accession |
|---|---|---|---|
| Liver Cirrhosis | Cirrhosis | Qin, N. et al. [1] | ENA database, accession number ERP005860 |
| Colorectal Cancer | Colorectal | Zeller, G. et al. [2] | ENA database, accession number ERP005534 |
| Inflammatory Bowel Disease | IBD | Qin, J. et al. [3] | EBI database, accession number ERA000116 |
| Obesity | Obesity | Le Chatelier, E. et al. [4] | ENA database, accession number ERP003612 |
| Type 2 Diabetes | WT2D (European Women Cohort) | Karlsson, F. H. et al. [5] | SRA database, accession number ERP002469 |
| | T2D (Chinese Cohort) | Qin, J. et al. [6] | SRA database, accession numbers SRA045646 and SRA050230 |

Table S1: Distribution of samples within the datasets of reference

The contents of the 'S1-Datasets' folder are as follows:

- **Datasets:** All original dataset files, in txt format ('abundance_[disease].txt'). These files contain the relative abundances of all OTUs sequenced from the original reads, and described by their precise taxonomy. Metadata per sample include identifiers for samples and subjects, disease state, age, BMI, country and gender of the subject the sample was taken from, the sequencing technology used for this sample and the subsequent analysis pipeline used to obtain OTU abundances.

- **Extracted Label files:** Contains vector files describing, respectfully of the original dataset's order, each individual's state in relation to the disease (i.e: 0 if control, 1 if unhealthy). They are directly extracted from the previously described original datasets, and formatted to be valid inputs for our implementation of the SPARTA pipeline.

### 3 The SPARTA pipeline

Unit tests for all of the major steps of the SPARTA pipeline are available in the S2 folder. Detailed instructions to run the unit tests and check the validity of results are given in the included 'testing_instructions.txt' file, and a python script for automatic verification of the outputs is also included. A description of each step, and a detailed justification of their expected outputs are given in the 'manual_verification.txt' file.

A full implementation of the pipeline is available in the 'SPARTA' folder, and on github at `https://github.com/baptisteruiz/SPARTA`.

### 4 Results

#### 4.1 SPARTA outputs per run

All of the outputs obtained per repetition of SPARTA, for all datasets, are available in the S3 folder.

The contents are organized as follows:

- **Intermediary outputs:** This archive includes files that are only intermediary in SPARTA's process and not listed as main outputs, but remain nonetheless relevant to the discussion and for reproducing the article's figures or manipulations. These include:
  - **Abundance per profile:** For each dataset, the abundance profiles of annotations and OTUs, restricted to the samples of a specific profile ('Sick' or 'Control') are given. Files are named as such: *[dataset_name]_tfigmavg_[count or score]_[sick or ctrl].csv.* Files labeled with 'count' contain information about the OTUs, those with 'score' pertain to the functional annotations.
  - **EsMeCaTa out:** For each dataset, the EsMeCaTa outputs relevant to SPARTA. This information is available in the following original nomenclature: '[dataset_name]/esmecata_outputs_annots/annotation_referenc
    Each OTU, numbered following the pipeline's internal nomenclature (see OTU name references folder), has a tsv tabular file named after it. This file contains the details of the OTU's meta-proteome, with the names and identifiers of all proteins, the names of the expressed genes, and the list of functional annotations associated to each protein (GO terms, EC numbers).
  - **OTU name references:** Per dataset, a tsv tabular file describes the pipeline's identifiers for OTUs, under the format 'OTU_X' with X being a number, and the detailed taxonomic affiliation of the original. These files are formatted to be used as input for the EsMeCaTa pipeline.
  - **OTU name references:** Per dataset, a csv tabular describes which samples were selected to be part of the test sets, per run.
- **Performance tables:** In a folder per SPARTA run applied to each dataset, the details of the effectuated classifications are compiled in the following files:
  - 'Performance_dataframe_entree_DeepMicro_[dataset]_[iteration number]_[tfigm]_[OTU].csv': These tabulars detail, for each Random Forest model trained in this ieration of the concerned SPARTA run, the optimal found training parameters, the optimal probability threshold for classification, and the classification performances (ROC AUC) on the training, validation and test subsets. On some earlier runs, the details of the indices of samples in the validation subset were given, but that information is not a constant. Concerning the files' nomenclature:

iteration number : does not appear for the files concerning the first iterations tfigm only appears for functional profiles scaled with the TF-IGM method. As this became the norm for our manipulations, later iterations do not necessarily contain information about non TF-IGM treated functional datasets.

OTU only appears if the file concerns an OTU abundance profile. Functional profiles are not specified.

- 'data_recap_[run number]_[dataset]_[tfigm].csv': these tabulars give the results of tests conducted on the distributions of the OTU and functional (TF-IGM treated or not) performances. For each profile, the maximum average performance and the selection iteration at which it was obtained are given. If applicable, the best distribution is compared to the first through a Mann-Whitney u-test, or in the case of the first runs, the significance of the evolution of the performance distributions is tested through a Kruskal-Wallis test. The top OTU ad annotation distributions are then compared through a Mann-Whitney u-test. The p-values of all tests are given.
- 'perf_plot_[dataset]_[tfigm].png': a visualization of the obtained results, showing violinplots af all classification performance distributions obtained on OTU profiles, and functional profiles (TF-IGM scaled or not, depending on the file's nomenclature).

- **Raw SPARTA outputs:** These are the detailed, non-compiled outputs of each SPARTA run conducted on each dataset, with or without TF-IGM normalization. The outputs are organized as follows:
  - **Classification performances :**
    - 'Data_leak_free_[dataset]_performances.txt': Performance scores (AUC) of the best performing RF model on the independent test functional datasets
    - 'Data_leak_free_[dataset]_OTU_performances.txt': Performance scores (AUC) of the best performing RF model on the independent test OTU datasets
    - run i (for all i iterations of the method):
      - 'OTU_classif_perfs.txt': Raw performance metrics of each RF model trained during this iteration on the internal verification subset of the OTU data (In order of the columns: Classification method, ROC AUC, Accuracy, Recall, Precision and F1 score).
      - 'SoFA_classif_perfs.txt': Raw performance metrics of each RF model trained during this iteration on the internal verification subset of the functional data (In order of the columns: Classification method, ROC AUC, Accuracy, Recall, Precision and F1 score).
  - **Selection outputs:**
    - run i (for all i iterations of the method)
      - OTU_[dataset].csv : List of all OTUs ranked by decreasing importance, with the importance cutoff integrated.
      - scores_[dataset].csv : List of all functional annotations ranked by decreasing importance, with the importance cutoff integrated (also available in Excel format).
  - **SoFAs**
    - OTU_table_stripped.tsv : Original OTU count table with new OTU names and without metadata.
    - SoFA_table.tsv : functional score profiles calculated for each sample.

#### 4.2 Aggregated selection outputs from SPARTA and DESeq2

The information gathered from the aggregation of the repeated selections by SPARTA and DESEq2, made in the context of the article, is available in folder S4. Contents are as follows:

- **Full Core Meta sublists:** Per iterative selection level of the pipeline (referred to as 'runs' in this instance) for each dataset, the lists of robust (Core) and candidate (meta) OTUs and annotations from SPARTA's automatic selection process are available under the names '[core_meta]_otus_[dataset]_tf_igm_run_[iteratio number].csv'. For the Meta outputs, the number of repetitions that selected a variable is given in the 'Count' column. For each selection level, Core and Meta sublists obtained from selecting the top X variables are available under the names '[core_meta]_otus_[dataset]_tf_igm_top_[X]_run_[iteration number].csv'. The Core and Meta sublists obtained from 10 applications of DESeq2 to the original dataset, with different p-value thresholds ($\alpha$) are available in the 'run 1' folders, under the na me 'core_annots_[dataset]_tf_igm_deseq_alpha_[$\alpha$]_run_run_1.csv'.

- **Full selection lists deseq:** This folder contains the full lists of selected variables by each application of DESeq2, at different selection thresholds, to the original OTU and functional datasets. These lists are collected, per dataset, in the 'OTU' (for OTU profiles) and 'SoFA' (for functional profiles) folders, under the names 'deseq_[$\alpha$]_selection_Run_[run number]'.

- **Aggregated selection information:** This folder contains overviews of the sizes of each selection, obtained by each method on the different profiles. The information is stored in csv tabulars under the nomenclature: 'Resume_[dataset]_[OTU/SoFA].csv'.

### 5 Reproduction of figures

The 'Code for figures' folder contains notebook codes that can be used, in the context of the given folders, to recreate the figures given in the article.

The following environments and packages are required for the code:

- **Fig 2:** Python environment with libraries seaborn, pandas, numpy, matplotlib and scipy.
- **Fig 3:** Python environment with libraries seaborn, pandas, numpy and matplotlib.
- **Fig 4:** R environment with packages eulerr and dict.
- **Fig 5:** Python environment with libraries seaborn, pandas, numpy and matplotlib.
- **Fig 6:** Python environment with libraries seaborn, pandas, numpy, matplotlib, networkx, json and sklearn.

The notebooks gather information automatically and directly from the outputs in the other folders, if the archive is extracted as is. Figures 5 and 6 gather information presented differently, which is stored in the 'Formatted info for fig 5 6' folder. Its contents are:

- '[annot/otu]_plus_info.csv': contains the list of robust (Core) annotations or OTUs, coupled with information concerning its name, average importance, and associated counterparts.
- '[annot/otu]_non_signif_plus_info.csv': contains the list of non-robust annotations or OTUs, coupled with information concerning its name, average importance, and associated counterparts.
- 'annot_plus_info_avg_pres.csv': same as 'annot_plus_info.csv', with extra information pertaining to the average presence of the annotations in control and unhealthy profiles.

Some intermediary files for Fig 6 are also long to compute (approx. 2 to 3 hours). For faster results, users have the option to import pre-calculated values instead, stored in the 'saved calculated info for fig 6' folder.
