## Supplementary figures and images for "SPARTA: Interpretable functional classification of microbiomes and detection of hidden cumulative effects"

### Supplemental Figure S1

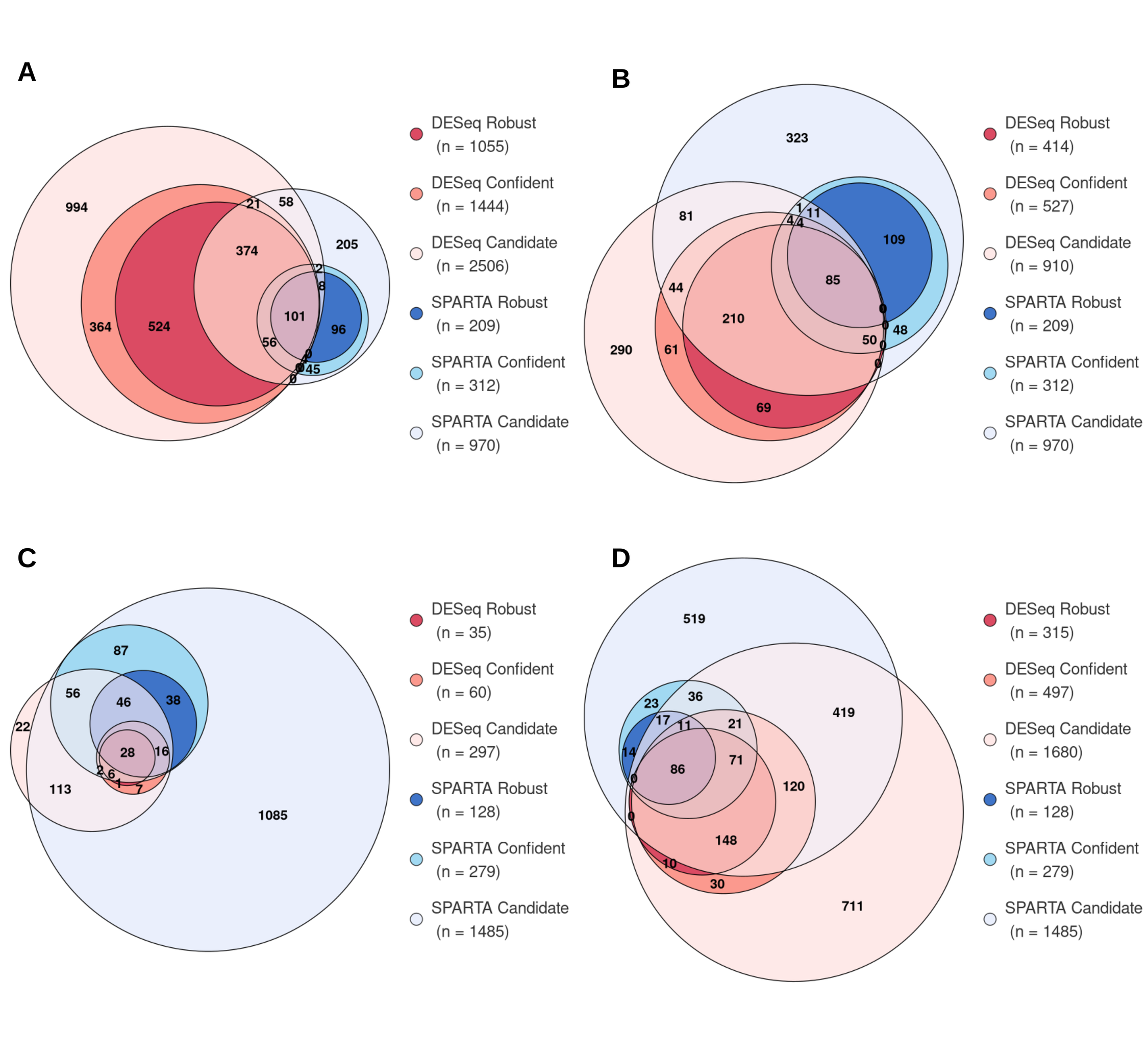

### Supplemental Figure S2

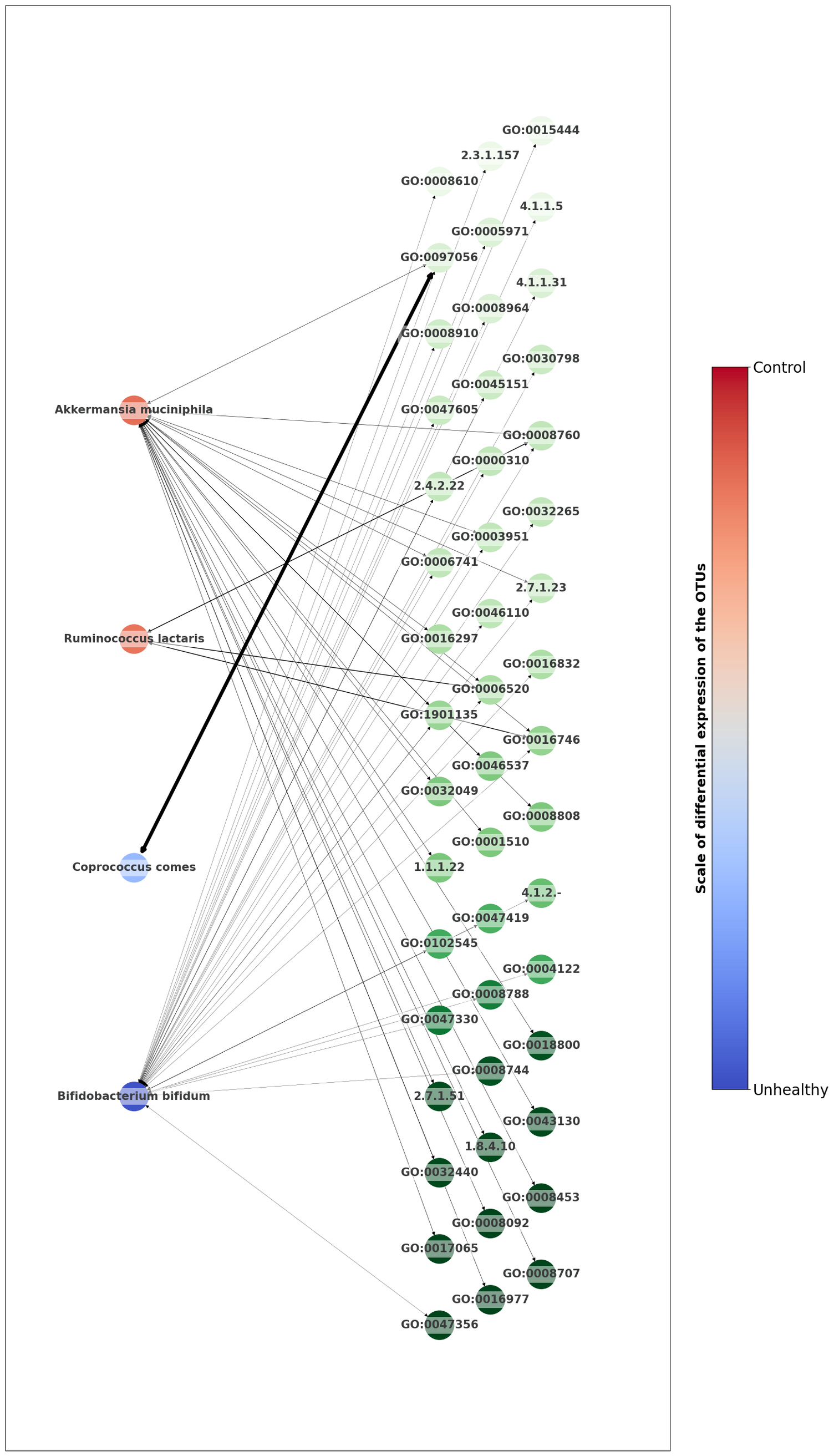

### Supplemental Table S1

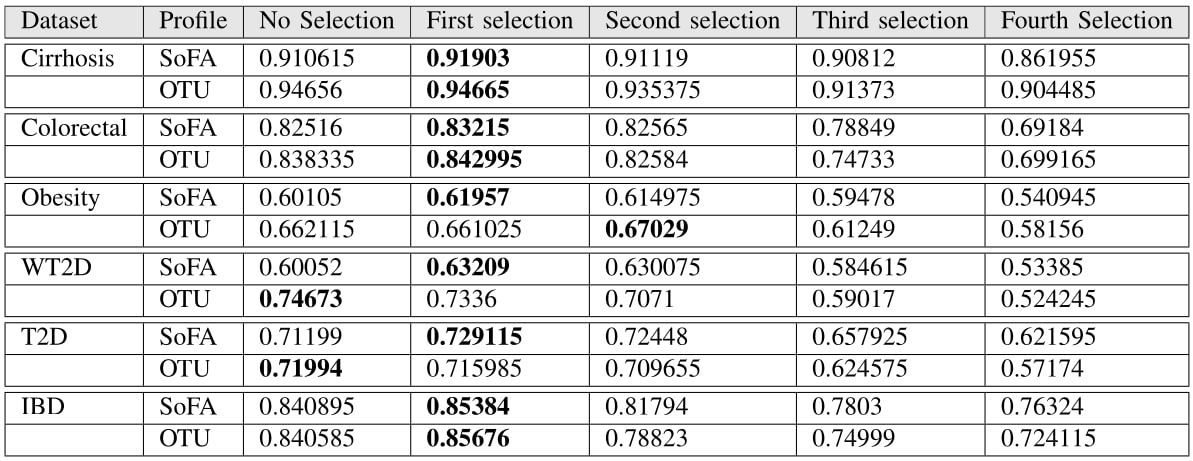

### Supplemental Table S2

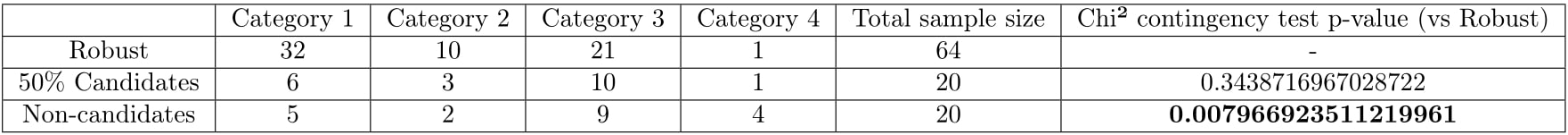
